# A new model for vertebrate mineralization via stabilized amorphous calcium carbonate for avian eggshell formation

**DOI:** 10.1101/2020.04.08.031989

**Authors:** Lilian Stapane, Nathalie Le Roy, Jacky Ezagal, Alejandro B. Rodriguez-Navarro, Valérie Labas, Lucie Combes-Soia, Maxwell T. Hincke, Joël Gautron

**Affiliations:** BOA, INRAE, Université de Tours, 37380 Nouzilly, France; Departmento de Mineralogia y Petrologia, Universidad de Granada, 18071 Granada, Spain; UMR PRC, INRAE 85, CNRS 7247, Université de Tours, IFCE, 37380 Nouzilly, France; Department of Innovation in Medical Education, and Department of Cellular and Molecular Medicine, University of Ottawa, Ottawa, Canada K1H8M5

**Keywords:** Biomineralization, Avian eggshell, extracellular vesicle, ACC transport, matrix proteins

## Abstract

Amorphous calcium carbonate (ACC) is an unstable mineral phase, which is progressively transformed into aragonite or calcite in biomineralization of marine invertebrate shells or avian eggshells, respectively. We have previously proposed a model of vesicular transport to provide stabilized ACC in chicken uterine fluid where mineralization takes place. Herein, we report further experimental evidence for this model. We confirmed the presence of extracellular vesicles (EVs) that contain ACC in uterine fluid using transmission electron microscopy and elemental analysis. We also demonstrate high levels of expression of vesicular markers in the oviduct segments where eggshell is formed. Moreover, proteomics and immunofluorescence confirmed the presence of major vesicular, mineralization-specific and eggshell matrix proteins in the uterus and in purified EVs. We propose a comprehensive role for EVs in eggshell mineralization, in which annexins transfer calcium into vesicles and carbonic anhydrase 4 catalyzes the formation of HCO_3_^−^, for accumulation of ACC in vesicles. We hypothesize that ACC is stabilized by ovalbumin and/or lysozyme or additional vesicle proteins identified in this study. Finally, EDIL3 and MFGE8 are proposed to serve as guidance molecules to target EVs to the mineralization site. We therefore report for the first time experimental evidence for the components of vesicular transport to supply ACC in vertebrate biomineralization. These results could give insight to understand the mineralization of otoconia, which are calcium carbonate biomineralized structures present in all vertebrates and necessary for balance and sensing linear acceleration.

---

Biomineralization is an extremely widespread process by which living organisms produce minerals; over 60 different biominerals have been identified, including calcium carbonates and phosphates (1). The formation of biominerals requires high local concentrations of calcium to form the crystalline polymorph. Recently, the transient amorphous mineral phase of calcium phosphates and carbonates has been recognized as a soluble and highly reactive source to facilitate the formation of the complex shapes of observed crystalline biomineral structures. The importance of amorphous calcium carbonate (ACC) as a transient mineral phase was reported in the calcification of calcium carbonate (CaCO_3_) biominerals (calcite, aragonite) (2–5). ACC is a metastable polymorph of CaCO_3_, and can provide high concentrations of ions for rapid physiological biomineralization (6). The high solubility of this transitory phase requires stabilization by regulatory molecules (2,5,7–11). Extracellular vesicles (EVs) were reported as candidates to play a role in the stabilization of ACC (5,9–11). Experimental evidence for EVs in ACC-mediated calcification in CaCO_3_ biominerals is only available for invertebrate structures such as sea urchin spines (calcite) and molluscan shells (2,12,13).

Amongst vertebrates, the avian class (Aves) appeared around 91 million years ago and is divided into Paleognathae (ancient birds such as ostrich, rhea and emu) and Neognathae (modern birds such as chicken, turkey or zebra finch) (14). All birds produce eggs with hard shells composed of calcitic calcium carbonate, which provides a biomineral barrier that allows appropriate development of the embryo within an autonomous chamber (15). In addition to defense against physical aggression, the eggshell protects the egg contents against microbial contamination, regulates water and gaseous exchange, and provides calcium for embryonic bone calcification (16,17). The chicken eggshell is a widely utilized experimental model. Eggshell formation occurs in the uterine segment of the hen oviduct and is one of the fastest processes of vertebrate biomineralization (18–20). Approximately 6 g of CaCO_3_ are deposited in a rapid period with a mineralization rate of 0.32 g/h (21). During this extracellular process, the uterine cells secrete organic and mineral eggshell precursors into the uterine fluid (UF) where mineralization takes place (22–24). Both mineral and organic precursors interact to produce the specific eggshell texture and its resulting mechanical properties (19,25). This process requires delivery of large amounts of calcium and carbonates at the site of eggshell calcification, which are continuously supplied from the blood across the uterine epithelium (26). The active transepithelial transfer of calcium and carbonates is well described and constitutes the current model for eggshell calcification (23,27).

In chicken eggshell, the important role of ACC during mineralization has been described (28). ACC aggregates are first massively deposited on the shell membranes and transform into calcite during initial calcification of the mammillary knobs (organic cores where the primary nucleation occurs). ACC is also required at the mineralization edge during the overall eggshell calcification process, in order to provide material for the growth of columnar calcite crystals that constitute the palisade layer (28). In a recent study, we used bioinformatics tools, gene expression and protein quantification to explore the role of EDIL3 and MFGE8 in chicken eggshell biomineralization. We hypothesized that EDIL3 and MFGE8 bind to EVs budding from uterine cells into the uterine fluid, in order to guide vesicular transport of stabilized ACC for delivery to the mineralizing site and moreover prevent non-specific precipitation (29).

In order to test this hypothesis, we used microscopic techniques to investigate exocytosis activity at the plasma membrane of uterine cells that could be a source of EVs in the uterine fluid. We also used elemental analysis to verify the presence of ACC inside these EVs and we quantified gene expression of a variety of validated EV components in the oviduct segments and other tissues. Finally, we purified EVs from UF and demonstrated the presence of key vesicular proteins in these vesicles. This experimental study is the first to demonstrate vesicular transport of ACC in vertebrates, which we propose supports the rapid eggshell biomineralization process in birds.

## Results

### Transmission electron microscopy of uterine epithelial cells and uterine fluid

The presence of vesicles in the tissues and milieu involved in shell mineralization was investigated by transmission electron microscopy (TEM). Ultra-thin negatively stained sections of uterus were examined by TEM to investigate exocytosis activity adjacent to the luminal site of mineralization (Fig. 1). Uterine ciliated cells possess numerous vacuolar and vesicular structures as well as dense and light granules (Fig. 1A). At higher magnification (Fig. 1B-D) we observed vesicles in the cell cytoplasm, their accumulation at the apical plasma membrane, and their budding to generate EVs in the adjacent luminal uterine fluid (Fig. 1C-D). Figure 1E and F, show EVs in the uterine fluid, with vesicle diameters in the 100-400 nm range. Vesicle membranes appeared electron-dense compared to their internal regions. This density is due to organic material comprising the membranes (glycoproteins, proteins, lipids…), which was stained by uranyl acetate. However, some internal regions also appeared electron-dense. These TEM observations demonstrated the presence of vesicles in the luminal uterine fluid adjacent to the apical region of uterine cells. The uterine epithelium contains ciliated and non-ciliated cells, and the same results were observed in proximity to non-ciliated cells (data not shown). UF was also subjected to TEM analysis (Fig. 2 and 3), where numerous EVs varying in diameter from 100 to 500 nm were observed (Fig. 2A, C and Fig. 3A).

**Figure 1.**
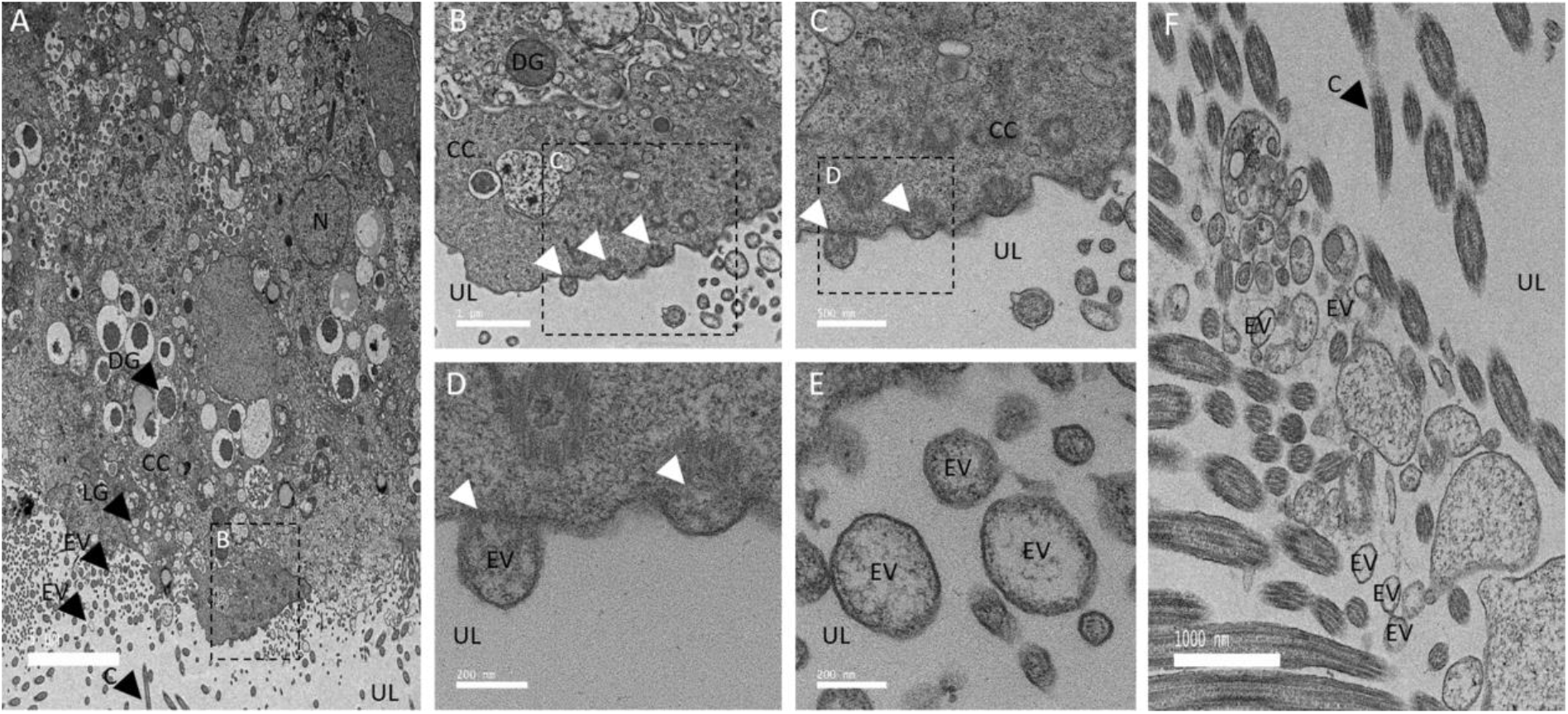
Transmission electron microscopy (TEM) of uterine cells. (A) TEM micrograph of uterine epithelium showing secretory activity (16 h p.o.). (B, C, D) Higher magnification TEM micrographs of uterine cells showing examples of exocytosis activity. (E, F) TEM micrographs illustrating the presence of numerous EVs in the uterine lumen (UL) at 7 h (F) and 16 h p.o. (E). Thin sections of uterus were negatively stained with 2 % uranyl acetate. C: cilia, CC: ciliated cells, DG: dense granule, LG: light granule, EV: extracellular vesicle, N: nucleus, UL: uterine lumen. White arrowheads indicate vesicle budding. Bars: A= 5 μm, B and F= 1 μm, C= 500 nm, D and E= 200 nm.

**Figure 2.**
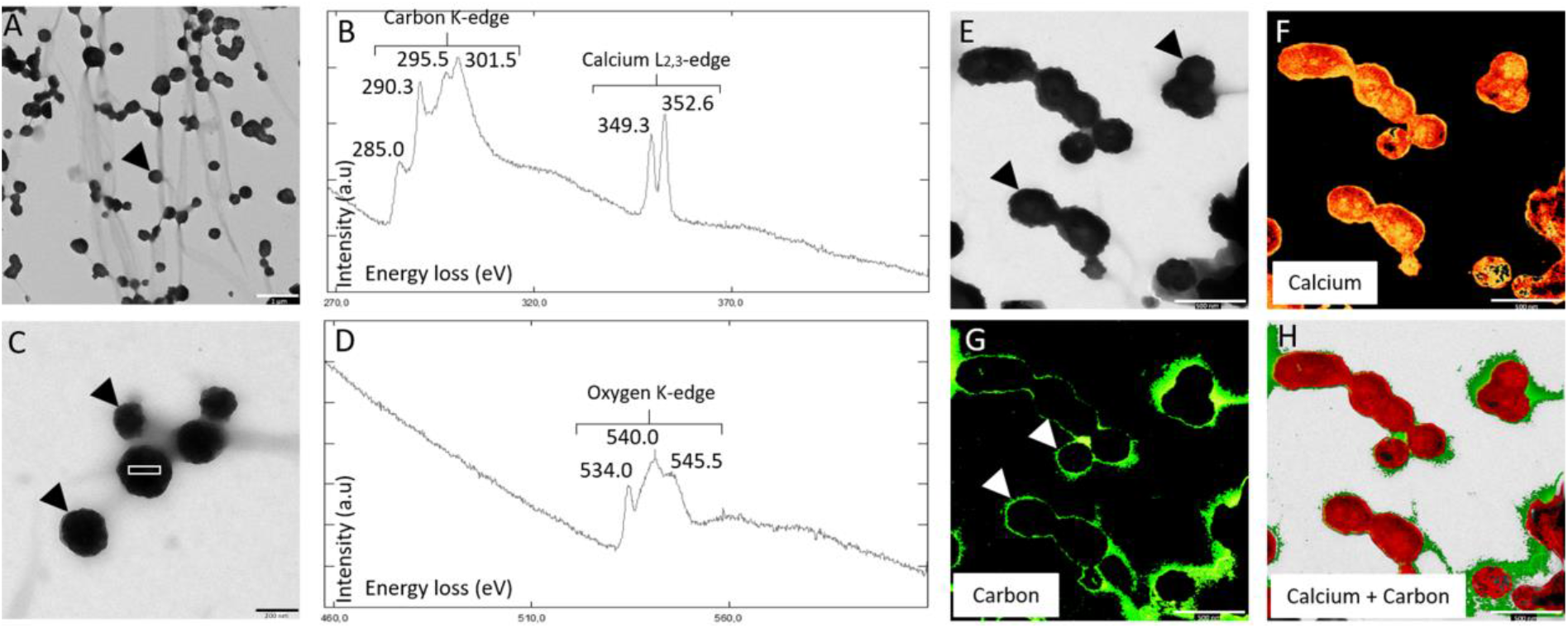
Transmission electron microscopy (TEM) on uterine fluid (UF) and elemental analysis (EELS) (A, C) TEM micrograph of EVs observed in UF. Black arrowheads indicate the EVs. White boxes indicate EELS analysis area of (B) and (D) graphs. (B, D) EELS spectroscopy on EVs from UF. The carbon K-edge shows three major peaks at 290.3, 295.5 and 301.5 electron volts (eV), characteristic of CaCO_3_. The peak at 285.0 from the carbon K-edge reflects the presence of organic material. The two peaks 349.3 and 352.6 eV define the calcium L2,3-edge of EELS. The oxygen K-edge displays a major peak at 540.0 specific to the carbonate group (CO_3_^2−^) and two other peaks at 534.0 and 545.5 indicate C=O bond. (E) TEM micrograph of EV, observed in UF with associated mapping of (F) calcium element (324-355 eV). (G) Organic phase carbon (280-290 eV) and (H) calcium + carbon combined elements detected by the EELS analysis. Bars A = 1 μm, C = 200 nm, E to H = 500 nm.

**Figure 3.**
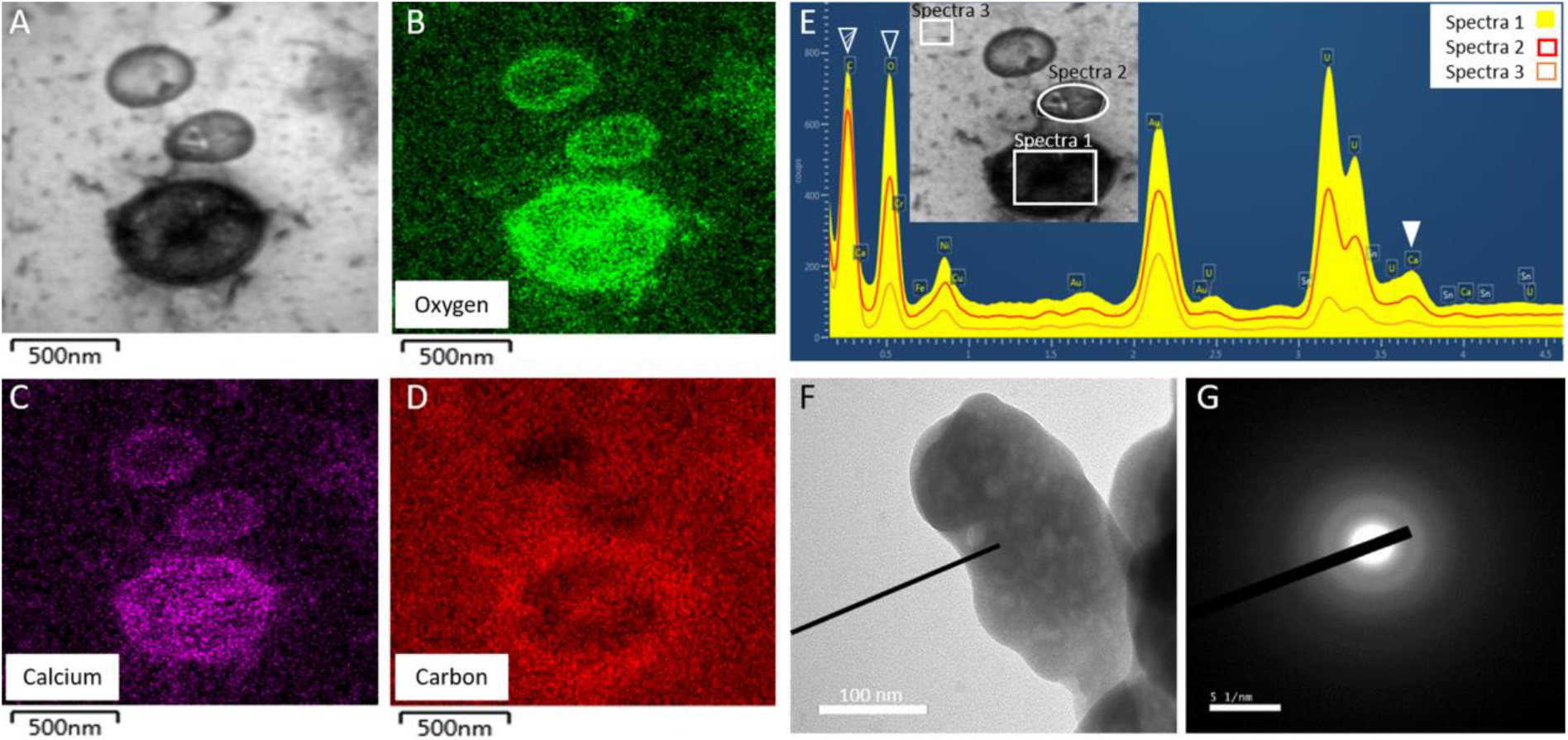
Transmission electron microscopy (TEM) on uterine fluid, energy-dispersive X-ray spectroscopy (EDS) and selected area electron diffraction (SAED) analyses. (A) Transmission electron microscopy micrograph (TEM) of EVs observed in uterine fluid (UF) fraction at 16 h p.o. with associated mapping of (B) oxygen element; (C) calcium element; and (D) carbon element; as detected by the energy-dispersive X-ray spectroscopy (EDS) analysis. (E) EDS spectrum of the EVs observed in UF. Filled white arrowhead indicates the calcium peak (3.690 keV) of EV whereas striped and empty white arrowheads show carbon (0.277 keV) and oxygen (0.525 keV) peaks, respectively. Boxes point out the areas of EDS analysis. (F, G) selected area electron diffraction (SAED) pattern of the mineral detected in EV of UF. Diffuse rings detected are characteristic of the ACC phase. Bars A to D = 500 nm, F = 1 μm, G = 5 1±nm.

### Elemental analysis to study chemical components of extracellular vesicles

At collection, uterine fluid was immediately frozen in liquid nitrogen in order to preserve EVs and their cargo. The elemental composition of the vesicular constituents was examined using electron energy loss spectroscopy (EELS, Fig. 2), and energy-dispersive X-ray spectroscopy (EDS, Fig. 3).

Using EELS, peaks from the carbon-K-edge (285.0 eV, 290.3 eV, 295.5 eV and 301.5 eV) and calcium L-2, 3-edge (349.3 eV and 352.6 eV) were detected (Fig. 2B, Table S1). All peaks, except 285.0 eV, are specific to CaCO_3_ (30). The analysis also revealed a major oxygen K-edge peak at 540.0 eV, which is also specific to the carbonate CO_3_^2−^ group (Fig. 2D, Table S1) (30). The energy loss graphs therefore revealed an elemental pattern specific to CaCO_3_ mineral (28,31). Moreover, the 285.0 eV peak from the carbon K-edge indicated the presence of the C=C bond and the two peaks from the oxygen K-edge (534.0 eV and 545.5 eV) are specific to the C=O bond (30). These data therefore confirm the presence of an organic phase (phospholipids and proteins) as well as calcium carbonate in the uterine EVs. These data were extended by elemental analysis with mapping of the carbon (237-290 eV) and calcium (324-355 eV) signals (Fig. 2E-H). These results showed a calcium signal inside the EVs (Fig. 2F), whereas a carbon signal, specific to the organic phase (C=C), was visualized at the vesicle periphery (Fig. 2G). Merger of the carbon and calcium maps (Fig. 2H) clearly shows that the EV membranes (phospholipid and proteins) enclose the calcium signal.

The EELS results were confirmed using energy-dispersive X-ray spectroscopic (EDS) analysis on the uterine fluid EVs (Fig. 3). Oxygen, carbon and calcium elements co-localized within EVs (Fig. 3A-D). A high background signal is notable for carbon because of the use of carbon grids. The difference between EDS spectrum 1 (EV), spectrum 2 (EV) and spectrum 3 (background) confirmed the presence of significant calcium and oxygen amounts inside EVs, compared to the background signal (Fig. 3E). Indeed, calcium and oxygen signals displayed 3- and 5-fold increases in EV locations (spectrum 1), compared to the background signal (spectrum 3), respectively. The question arises regarding the form of calcium carbonate inside the vesicles. In order to identify the mineral phase within EVs, we carried out selected area electron diffraction (SAED) on EVs observed in UF (Fig. 3F, G). We detected diffuse continuous rings indicating that the calcium carbonate present inside vesicles was amorphous, similar in nature to the ACC identified during eggshell formation (28).

### Vesicular gene expression in various Gallus gallus tissues

We evaluated the literature on bone and cartilage extracellular vesicles (32–35), EVs (36–39), and the Vesiclepedia database (www.microvesicles.org/), which lists the top 100 proteins that are identified in EVs (40). A total of 33 genes coding for proteins involved in vesicular transport were selected, corresponding to proteins involved as calcium channels to supply calcium, bicarbonate supplier/transporter, chaperone molecules, addressing molecules, intracellular trafficking proteins, extracellular biogenesis and release, and signaling proteins (Table S2). Expression levels for these 33 genes were quantified in different tissues and organs, namely oviduct segments, bone, duodenum, kidney and liver (Table 1, Fig. 4A). The oviduct is the reproductive tract, where the forming egg transits as it progressively acquires specific egg compartments. The egg white components and eggshell membrane precursors are secreted and deposited in the magnum (Ma) and white isthmus (WI) segments, respectively, while the red isthmus (RI) and uterus (Ut) are involved in the onset and the development of shell mineralization. Tissue samples from these four specialized oviduct regions were collected to evaluate expression of EV markers associated with egg white deposition (Ma), eggshell membrane formation (WI), and shell calcification (RI, Ut). Bone was selected as a mineralized tissue (hydroxyapatite) where EVs have been demonstrated (32,33,41,42). Duodenum (D) and kidney (K) exhibit active ion transport activity without any associated calcification. Finally, liver (L) was selected as an important organ involved in general metabolism. Comparisons of quantified gene expression in these organs and tissues were displayed using a heatmap diagram (Fig. 4A). Z-scores are expressed in terms of standard deviations from their means for each gene. Consequently, the color indicates an intuitive idea of the relative variation of each gene in the different tissues. We observed the highest Z-scores in oviduct segments (Ma, WI, RI, Ut), for 20 vesicular genes (*Anxa1*, *Anxa2*, *Anxa8, Ap1g1*, *Cd9, Cd82, Edil3, Hspa8, Itgb1, Pdcd6ip, Rab5a, Rab27a, Sdcbp, Tsg101, Vamp3, Vamp7, Vps4, Vps26a, Ywhah* and *Ywhaz)* compared to the other tissues (Clusters 6 to 8, Fig. 4A). Expression of genes was also analyzed using ANOVA and Tukey pairwise analysis (Table 1). With the notable exception of *Ap1g1*, all other genes with highest Z-scores in uterus were also significantly different in the same uterine tissue using ANOVA and pairwise analysis (Table 1). Additionally, these statistical tests show that *Itgb1, Sdcbp, Vamp7* and *Vps4b* were also significantly over-expressed in one or several other tissues (bone (B), kidney (K), and duodenum (D)). *Anxa5, Anxa11, Ap1g1, Hsp90b* and *Rab7a* were not differentially expressed in the various tissues tested. Both *Rab11a* and *Slc4a7* were significantly over-expressed in D, while *Ca2* and *Anxa7* were significantly over-expressed in B and K respectively to other tissues. *Arf6* exhibited a significant higher expression in D and uterus. In this test, different letters are used for statically significant different level of expression. Analysis of the remaining three genes *Anxa6, Ralb,* and *Vcp* reported most of commons letters indicating not significantly different expression for these genes (Table 1).

**Table 1.** Normalized vesicular gene expression in the four oviduct regions and other tissues. Expression results are reported as mean + SD. Gene expression was assessed during the active growth phase of eggshell mineralization at 10 h p.o. except for B at 18 h p.o.. Following reverse transcription, gene expression was quantified using primers designed with Primer-blast tool of NCBI (https://www.ncbi.nlm.nih.gov/tools/primer-blast/) and validated for the Biomark microfluidic system (BMK-M-96.96; Fluidigm). Relative quantification was normalized with eight housekeeping genes using GenNorm software. Superscript letters “a” to “e” correspond to significance results of the ANOVA-Tukey tests with “a” the letter corresponding to the highest level of expression for each gene. Tibial bone, B; duodenum, D; kidney, K; liver, L; magnum, Ma; white isthmus, WI; red isthmus, RI; uterus, Ut. The gene accession numbers and primer sequences are compiled in Table S4.

| Tissues |  |  |  |  |  | Oviduct segments |  |  |  | ANOVA<br>p-value |
| --- | --- | --- | --- | --- | --- | --- | --- | --- | --- | --- |
| Function | Gene | B | D | K | L | Ma | WI | RI | Ut |  |
| Calcium<br>channel | *Anxa1* | 0.205±0.200<sup>cd</sup> | 0.002±0.001<sup>d</sup> | 0.009±0.004<sup>d</sup> | 0.001±0.001<sup>d</sup> | 0.194±0.066<sup>cd</sup> | 2.190±0.749<sup>b</sup> | 3.338±0.768<sup>a</sup> | 0.891±0.320<sup>c</sup> | <0.001 |
| | *Anxa2* | 0.307±0.240<sup>cd</sup> | 0.521±0.180<sup>cd</sup> | 0.006±0.009<sup>d</sup> | 0.001±0.002<sup>d</sup> | 0.543±0.340<sup>cd</sup> | 1.208±0.805<sup>bc</sup> | 2.884±1.247<sup>a</sup> | 1.904±0.459<sup>ab</sup> | <0.001 |
| | *Anxa5* | 0.431±0.359 | 0.401±0.289 | 0.189±0.173 | 1.462±0.828 | 2.001±2.920 | 0.560±0.298 | 1.405±1.044 | 0.298±0.193 | 0.068 |
| | *Anxa6* | 2.116±1.522<sup>ab</sup> | 0.446±0.282<sup>b</sup> | 0.313±0.163<sup>b</sup> | 1.473±0.558<sup>ab</sup> | 1.328±1.140<sup>ab</sup> | 0.478±0.375<sup>ab</sup> | 0.347±0.147<sup>b</sup> | 0.543±0.437<sup>ab</sup> | 0.008 |
| | *Anxa7* | 0.206±0.081<sup>cd</sup> | 1.436±0.306<sup>b</sup> | 7.074±1.170<sup>a</sup> | 0.191±0.059<sup>d</sup> | 0.623±0.330<sup>bcd</sup> | 0.676±0.410<sup>bcd</sup> | 1.122±0.284<sup>bc</sup> | 1.450±0.526<sup>b</sup> | <0.001 |
| | *Anxa8* | 0.026±0.029<sup>d</sup> | 0.001±0.001<sup>d</sup> | 0.001±0.001<sup>d</sup> | 0.001±0.001<sup>d</sup> | 1.669±0.922<sup>bc</sup> | 2.513±1.205<sup>ab</sup> | 3.958±1.720<sup>a</sup> | 0.979±0.344<sup>cd</sup> | <0.001 |
| | *Anxa11* | 0.214±0.044 | 0.562±0.152 | 0.644±0.293 | 0.916±0.939 | 3.760±6.150 | 2.180±3.080 | 1.615±1.436 | 1.008±0.647 | 0.236 |
| Bicarbonate<br>supplier/<br>transporter | *Ca2* | 21.440±10.270<sup>a</sup> | 2.585±1.127<sup>b</sup> | 1.633±0.834<sup>b</sup> | 0.512±0.157<sup>b</sup> | 0.763±0.196<sup>b</sup> | 1.149±0.297<sup>b</sup> | 0.561±0.171<sup>b</sup> | 1.131±0.394<sup>b</sup> | <0.001 |
| | *Slc4a7* | 0.115±0.039<sup>cd</sup> | 1.905±0.665<sup>a</sup> | 0.069±0.008<sup>cd</sup> | 0.012±0.007<sup>d</sup> | 0.454±0.149<sup>cd</sup> | 0.551±0.287<sup>c</sup> | 1.078±0.262<sup>b</sup> | 0.529±0.120<sup>c</sup> | <0.001 |
| Membrane<br>organizer | *Cd82* | 0.634±0.345<sup>cd</sup> | 0.769±0.310<sup>bcd</sup> | 0.367±0.241<sup>cd</sup> | 0.162±0.141<sup>d</sup> | 1.598±0.970<sup>ab</sup> | 0.943±0.260<sup>abcd</sup> | 1.270±0.403<sup>abc</sup> | 1.865±0.834<sup>a</sup> | <0.001 |
| | *Cd9* | 0.085±0.100<sup>b</sup> | 0.258±0.129<sup>b</sup> | 0.403±0.168<sup>b</sup> | 0.053±0.068<sup>b</sup> | 1.751±1.255<sup>b</sup> | 12.900±4.98<sup>a</sup> | 1.780±0.323<sup>b</sup> | 1.617±1.106<sup>b</sup> | <0.001 |
| Chaperone | *Hsp90b1* | 2.427±1.776<sup>a</sup> | 0.941±0.672<sup>a</sup> | 3.230±3.500<sup>a</sup> | 5.250±3.920<sup>a</sup> | 4.560±4.650<sup>a</sup> | 1.649±0.760<sup>a</sup> | 1.627±0.659<sup>a</sup> | 2.992±0.840<sup>a</sup> | 0.039 |
| | *Hspa8* | 0.562±0.125<sup>d</sup> | 0.840±0.203<sup>bcd</sup> | 1.180±0.334<sup>b</sup> | 0.404±0.151<sup>d</sup> | 0.973±0.249<sup>bc</sup> | 0.919±0.287<sup>bc</sup> | 1.709±0.399<sup>a</sup> | 1.084±0.205<sup>b</sup> | <0.001 |
| Adhesion<br>molecule | *Itgb1* | 1.925±0.947<sup>a</sup> | 0.345±0.084<sup>bc</sup> | 0.671±0.491<sup>abc</sup> | 0.133±0.064<sup>c</sup> | 1.827±0.961<sup>a</sup> | 1.536±0.826<sup>ab</sup> | 1.783±1.054<sup>a</sup> | 1.396±0.504<sup>abc</sup> | <0.001 |
| | *Edil3* | 0.019±0.017<sup>b</sup> | 0.000±0.000<sup>b</sup> | 0.001±0.003<sup>b</sup> | 0.000±0.000<sup>b</sup> | 0.000±0.000<sup>b</sup> | 0.825±0.303<sup>a</sup> | 1.101±0.737<sup>a</sup> | 0.737±0.464<sup>a</sup> | <0.001 |
| | *Mfg8* | 0.514±0.380<sup>c</sup> | 0.114±0.078<sup>c</sup> | 0.856±0.208<sup>c</sup> | 4.373±1.223<sup>a</sup> | 0.029±0.063<sup>c</sup> | 0.803±0.399<sup>c</sup> | 2.139±1.194<sup>b</sup> | 0.381±0.385<sup>c</sup> | <0.001 |
| Intracellular<br>trafficking | *Rab11a* | 0.399±0.125<sup>d</sup> | 2.575±0.600<sup>a</sup> | 0.993±0.110<sup>b</sup> | 0.506±0.079<sup>cd</sup> | 1.110±0.201<sup>b</sup> | 0.703±0.130<sup>bcd</sup> | 0.952±0.076<sup>bc</sup> | 1.116±0.226<sup>b</sup> | <0.001 |
| | *Rab27a* | 0.381±0.168<sup>c</sup> | 0.329±0.197<sup>c</sup> | 0.585±0.283<sup>c</sup> | 0.185±0.144<sup>c</sup> | 2.306±1.363<sup>a</sup> | 1.778±0.517<sup>ab</sup> | 1.090±0.485<sup>bc</sup> | 1.742±0.634<sup>ab</sup> | <0.001 |
|  | **Rab5a** | 0.525±0.353<sup>bc</sup> | 0.779±0.251<sup>bc</sup> | 0.784±0.324<sup>bc</sup> | 0.780±0.422<sup>bc</sup> | 1.701±0.881<sup>a</sup> | 1.337±0.500<sup>abc</sup> | 1.294±0.300<sup>abc</sup> | 1.649±0.596<sup>ab</sup> | **<0.001** |
|  | **Rab7a** | 1.879±0.849 | 1.399±0.781 | 0.991±0.381 | 1.016±0.500 | 1.375±0.523 | 1.258±0.335 | 1.353±0.377 | 1.509±0.354 | 0.297 |
|  | **Ralb** | 0.987±0.257<sup>ab</sup> | 1.034±0.658<sup>ab</sup> | 1.350±0.587<sup>a</sup> | 1.191±0.484<sup>ab</sup> | 0.679±0.939<sup>ab</sup> | 0.615±0.371<sup>ab</sup> | 0.644±0.438<sup>ab</sup> | 0.349±0.203<sup>b</sup> | 0.039 |
|  | **Vcp** | 1.116±0.495<sup>ab</sup> | 0.414±0.136<sup>b</sup> | 0.707±0.259<sup>b</sup> | 2.129±0.802<sup>a</sup> | 1.379±1.746<sup>ab</sup> | 1.071±0.350<sup>ab</sup> | 1.229±0.469<sup>ab</sup> | 1.334±0.519<sup>ab</sup> | 0.027 |
|  | **Arf6** | 0.597±0.141<sup>bc</sup> | 1.688±0.490<sup>a</sup> | 0.770±0.201<sup>bc</sup> | 0.702±0.360<sup>bc</sup> | 0.647±0.255<sup>bc</sup> | 0.453±0.194<sup>c</sup> | 0.949±0.305<sup>bc</sup> | 1.206±0.370<sup>ab</sup> | <0.001 |
| EV biogenesis | **Sdcbp** | 0.581±0.203<sup>d</sup> | 0.920±0.225<sup>cd</sup> | 2.398±0.326<sup>ab</sup> | 0.546±0.210<sup>d</sup> | 2.585±1.130<sup>a</sup> | 2.506±0.629<sup>ab</sup> | 2.013±0.386<sup>ab</sup> | 1.567±0.422<sup>bc</sup> | **<0.001** |
|  | **Tsg101** | 0.134±0.018<sup>e</sup> | 0.510±0.136<sup>cd</sup> | 0.764±0.180<sup>bc</sup> | 0.386±0.056<sup>de</sup> | 1.206±0.385<sup>a</sup> | 0.900±0.224<sup>ab</sup> | 1.088±0.134<sup>ab</sup> | 1.192±0.184<sup>a</sup> | **<0.001** |
|  | **Vps26a** | 0.447±0.271<sup>de</sup> | 0.831±0.186<sup>cde</sup> | 1.078±0.147<sup>bcd</sup> | 0.466±0.183<sup>de</sup> | 1.203±0.419<sup>abcd</sup> | 1.754±0.338<sup>ab</sup> | 1.937±0.657<sup>a</sup> | 1.417±0.771<sup>abc</sup> | **<0.001** |
|  | **Pdcd6ip** | 0.213±0.047<sup>e</sup> | 1.150±0.225<sup>bc</sup> | 0.727±0.156<sup>cde</sup> | 0.493±0.109<sup>de</sup> | 0.777±0.332<sup>cd</sup> | 0.842±0.303<sup>cd</sup> | 1.454±0.301<sup>b</sup> | 2.030±0.545<sup>a</sup> | **<0.001** |
| EV release | **Vamp3** | 0.199±0.060<sup>d</sup> | 0.575±0.088<sup>bc</sup> | 0.365±0.124<sup>cd</sup> | 0.156±0.080<sup>d</sup> | 0.682±0.190<sup>b</sup> | 0.751±0.122<sup>b</sup> | 1.117±0.253<sup>a</sup> | 0.749±0.134<sup>b</sup> | **<0.001** |
|  | **Vamp7** | 0.439±0.224<sup>de</sup> | 0.590±0.281<sup>cde</sup> | 1.527±0.542<sup>ab</sup> | 1.088±0.258<sup>abcd</sup> | 0.911±0.258<sup>bcde</sup> | 1.198±0.242<sup>abc</sup> | 1.407±0.552<sup>ab</sup> | 1.704±0.373<sup>a</sup> | **<0.001** |
|  | **Vps4b** | 0.014±0.010<sup>b</sup> | 0.041±0.020<sup>ab</sup> | 0.019±0.019<sup>b</sup> | 0.009±0.006<sup>b</sup> | 0.052±0.029<sup>ab</sup> | 0.028±0.022<sup>b</sup> | 0.029±0.017<sup>b</sup> | 0.090±0.069<sup>a</sup> | **<0.001** |
|  | **Ap1g1** | 0.613±0.163<sup>ab</sup> | 0.649±0.154<sup>ab</sup> | 0.509±0.097<sup>ab</sup> | 0.334±0.185<sup>b</sup> | 0.824±0.539<sup>ab</sup> | 0.726±0.404<sup>ab</sup> | 1.063±0.260<sup>ab</sup> | 1.199±0.465<sup>ab</sup> | 0.030 |
| Signaling protein | **Ywhah** | 0.338±0.089<sup>abcd</sup> | 0.517±0.310<sup>abcd</sup> | 0.171±0.104<sup>cd</sup> | 0.111±0.098<sup>d</sup> | 0.970±0.339<sup>ab</sup> | 1.071±0.683<sup>ab</sup> | 1.089±0.562<sup>a</sup> | 0.872±0.568<sup>abc</sup> | **<0.001** |
|  | **Ywhaz** | 0.485±0.200<sup>cd</sup> | 0.501±0.152<sup>cd</sup> | 0.507±0.131<sup>cd</sup> | 0.136±0.050<sup>d</sup> | 0.660±0.266<sup>c</sup> | 0.786±0.159<sup>bc</sup> | 1.192±0.291<sup>b</sup> | 2.121±0.394<sup>a</sup> | **<0.001** |

**Figure 4.**
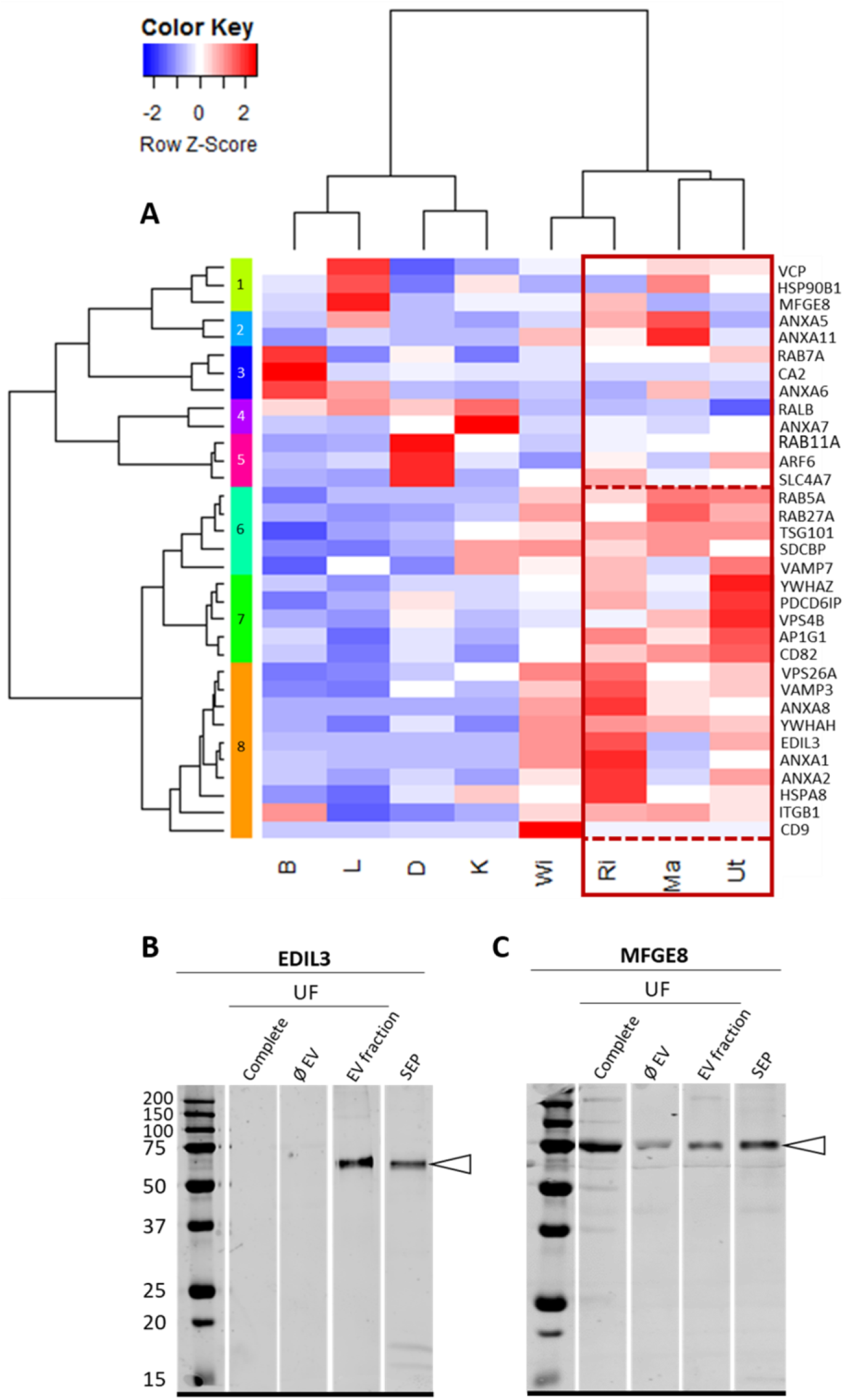
Heatmap of vesicular gene expression and Western blot analysis of EDIL3 and MFGE8. (A) Heatmap of vesicular gene expression in the four oviduct segments (Ma, magnum; WI, white isthmus; RI, red isthmus and Ut, uterus) and four tissues (B, bone; D, duodenum; K, kidney; L, liver). Z-score range was colored from blue (−2 Row Z-Score, low expression) to white (0 Row Z-Score, intermediate expression) and to red (2 Row Z-score, high expression). Gene accessions and primers are reported in Table S4. (B-C) Western blot analysis of (B) EDIL3 and (C) MFGE8 in uterine fluid (UF): unfractioned UF (complete), depleted UF (ØEV), EV fraction of UF (EV fraction) and soluble eggshell proteins (SEP). Proteins (15 μg) were subjected to 12.5 % SDS-page electrophoresis and blotted for analysis. The membranes were probed with (B) rabbit polyclonal anti-EDIL3 (1: 1,000, SAB2105802, Sigma-Aldrich, Saint-Quentin Fallavier, France) and (C) rabbit synthetic anti-MFGE8 peptide (1: 1,000, rabbit polyclonal; ProteoGenix, Schiltigheim). White arrowheads indicate the immunoreactive bands.

### Proteomic and Western blot analyses of EVs

To determine the protein composition of EVs isolated from the UF, we carried out proteomics analysis using nanoLC-MS/MS and verified these observations with Western blotting techniques. Twenty-nine non-redundant proteins were identified in our proteomics analysis (Table 2 and Table S3). Amongst them, seven proteins correspond to predicted major actors in the hypothetical vesicular transport of ACC (Carbonic Anhydrase-4 (CA4), EGF-like repeat and discoidin I-like domain-containing protein 3 (EDIL3), ezrin (EZR), programmed cell death 6-interacting protein (PDCD6IP), syntenin-1, ovalbumin (OVA) and lysozyme C (LYZ); Table 2 and 3) (29). In addition, several immunoglobulins and other proteins already identified in chicken eggshell matrix (ovotransferrin, ovalbumin-related X, ovalbumin-related Y, ovocalyxin-25, clusterin, ovomucin, vitelline membrane outer layer protein 1, ovostatin, ovoglobulinG2, prominin-1-A, alpha-2-macroglobulin, aminopeptidase N) were detected (Table 2). Finally, the angiotensin-converting enzyme precursor (ACE), was also identified.

**Table 2.** List of the 29 proteins identified by nanoLC-MS/MS analysis in uterine fluid EVs. The EV fraction was purified from uterine fluid (UF) collected at 9 h p.o.. Supplementary information on proteomic results are provided in Table S3. Indicated accessions correspond to the GenBank accession number.

| # | Identified Proteins | Symbols | Accession Number | GeneID | Theoretical molecular weight | Normalized weighted spectra | emPAI |
| --- | --- | --- | --- | --- | --- | --- | --- |
| 1 | Cluster of ovalbumin | OVA | AUD54558.1 | 396058 | 43 kDa | 1 031.10 | 29.909 |
| 2 | Cluster of ovotransferrin | OVOT | P02789.2 | 396241 | 78 kDa | 560 | 8.219 |
| 3 | Cluster of ovomucin | MUC | NP_989992.1 | 395381 | 234 kDa | 448.91 | 0.82205 |
| 4 | Cluster of alpha-2-macroglobulin-like | A2M | XP_025007673.1 | 418254 | 158 kDa | 288 | 0.89983 |
| 5 | Cluster of ovomacroglobulin (ovostatin) | OVST | CAA55385.1 | 396151 | 164 kDa | 215 | 0.47326 |
| 6 | Ig mu chain C region | N/A | P01875.2 | N/A | 48 kDa | 210 | 2.499 |
| 7 | immunoglobulin alpha heavy chain | N/A | AAB22614.2 | N/A | 62 kDa | 70 | 0.96212 |
| 8 | Ig light chain | N/A | AAA48859.1 | 416928 | 25 kDa | 63 | 0.88882 |
| 9 | ovalbumin-related Y | OVAL-Y | BAM13279.1 | 420897 | 44 kDa | 66 | 0.66086 |
| 10 | polymeric immunoglobulin receptor | PIGR | AAQ14493.1 | 419848 | 71 kDa | 71 | 0.31136 |
| 11 | ovoglobulinG2 | OVOG2/TENP | BAM13270.1 | 395882 | 47 kDa | 66 | 0.59817 |
| 12 | Cluster of immunoglobulin light chain variable region | N/A | AJQ23663.1 | N/A | 11 kDa | 42 | 0.87697 |
| 13 | Cluster of anti-prostate specific antigen antibody immunoglobulin variable region | N/A | ALX37966.1 | N/A | 13 kDa | 51 | 1.4661 |
| 14 | Cluster of programmed cell death 6-interacting protein | PDCD6IP | XP_015137020.1 | 420725 | 97 kDa | 46 | 0.16725 |
| 15 | Cluster of immunoglobulin light chain variable region | N/A | AJQ23661.1 | N/A | 11 kDa | 34 | 1.2195 |
| 16 | Cluster of Lysozyme C | LYZ | P00698.1 | 396218 | 16 kDa | 41 | 1.2327 |
| 17 | clusterin | CLU | NP_990231.1 | 395722 | 51 kDa | 7 | 0.36338 |
| 18 | Cluster of ovalbumin-related protein X | OVAL-X | NP_001263315.1 | 420898 | 44 kDa | 16 | 0.35247 |
| 19 | carbonic anhydrase 4 | CA4 | XP_415893.1 | 417647 | 36 kDa | 23 | 0.5467 |
| 20 | Cluster of ezrin | EZR | NP_990216.1 | 395701 | 69 kDa | 13 | 0.20218 |
| 21 | uncharacterized protein LOC771972 (OCX25) | OCX25 | XP_001235178.3 | 771972 | 27 kDa | 10 | 0.41449 |
| 22 | EGF-like repeat and discoidin I-like domain-containing protein 3 | EDIL3 | XP_424906.3 | 427326 | 54 kDa | 12 | 0.2673 |
| 23 | immunoglobulin heavy chain V-region | N/A | AAA48830.1 | N/A | 11 kDa | 9 | 1.2444 |
| 24 | Cluster of syntenin-1 | SDCBP | NP_001026195.1 | 421136 | 32 kDa | 11 | 0.21673 |
| 25 | aminopeptidase N | AMPN | ACZ95799.1 | 395667 | 109 kDa | 6 | 0.060894 |
| 26 | Cluster of immunoglobulin light chain variable region | N/A | AJQ23760.1 | N/A | 10 kDa | 6 | 0.75959 |
| 27 | prominin-1-A | PROM1 | XP_004942263.1 | 423849 | 88 kDa | 3 | 0.075475 |
| 28 | angiotensin-converting enzyme precursor | ACE | NP_001161204.1 | 419953 | 147 kDa | 5 | 0.044662 |
| 29 | vitelline membrane outer layer protein 1 | VMO1 | NP_001161233.1 | 418974 | 20 kDa | 4 | 0.35873 |

**Table 3.** Proteins of the proposed ACC extracellular vesicle system detected by nanoLC-MS/MS analysis (Scaffold software) in uterine fluid EVs. The EV fraction was purified from uterine fluid (UF) collected at 9 h p.o. The entire set of detected proteins is listed in Table S3. Indicated accessions correspond to the GenBank accession number. ACC, amorphous calcium carbonate; EV: extracellular vesicle; HCO_3_^−^, bicarbonate ion. Vesiclepedia database (www.microvesicles.org) classifies the proteins that are frequently identified in EVs. EV localization was predicted using molecular function of proteins on Uniprot (https://www.uniprot.org/). References: (32–35,62–65,67,69–78).

| Protein name | Accession number (GenBank) | Gene ID | Molecular weight | Hypothetical role in EV from UF | EV localization | Vesiclepedia rank | Bone/cartilage EV references | Eggshell/UF proteome references |
| --- | --- | --- | --- | --- | --- | --- | --- | --- |
| CA4 | XP_415893.1 | 417647 | 36 kDa | $\text{HCO}_3^-$ supplier | membrane: glycosylphosphatidylinositol (GPI-anchored) | N/A | N/A | 62, 67, 74, 76, 78 |
| EDIL3 | XP_424906.3 | 427326 | 54 kDa | targeting protein | membrane: phosphatidylserine-binding | N/A | 34 | 63, 65, 67, 70, 71, 72, 73, 75, 76, 78 |
| Ezrin | NP_990216.1 | 395701 | 69 kDa | membrane fusion (plasma membrane-cytoskeleton linker) | membrane: phosphatidylinositol 4,5 bisphosphate-binding | 30/100 | 33, 34, 35 | 62, 63, 74, 78 |
| PDCD6IP | XP_015137020.1 | 420725 | 97 kDa | EV biogenesis | lumen: incorporation | 1/100 | 32, 33 | 62, 77, 78 |
| Syntenin-1 | NP_001026195.1 | 421136 | 32 kDa | EV biogenesis | membrane: phosphatidylinositol 4,5 bisphosphate-binding | 19/100 | 32, 33, 34 | 62, 63, 64, 74, 75, 78 |
| Ovalbumin | AUD54558.1 | N/A | 43 kDa | ACC stabilization | lumen: incorporation | N/A | N/A | 62, 63, 65, 67, 69, 70, 74, 75, 76 |
| Lysozyme C | P00698.1 | 396218 | 16 kDa | ACC stabilization | lumen: incorporation | N/A | 34 | 62, 63, 65, 67, 69, 70, 74, 75, 76, 78 |

To confirm EDIL3 proteomic identification, we performed Western blot analysis on various samples: unfractioned UF (complete), EV fraction of UF (EV fraction), depleted UF (ØEV) and soluble eggshell proteins (SEP) (Fig. 4B, C). We also carried out immunoblotting for milk fat globule-EGF factor 8 (MFGE8), since it is another protein potentially involved in guiding EVs during shell mineralization (29). Immunoreactive bands in the vesicle fractions were observed using both antibodies (anti-EDIL3 and anti-MFGE8). Using anti-EDIL3 antibodies, a unique immunoband was observed around 60-kDa (54-kDa predicted molecular weight), only in the EV fraction and SEP (Fig. 4B). MFGE8 exhibited a single band around 75-kDa (53-kDa predicted molecular weight) in all samples (Fig. 4C). The higher than predicted molecular mass observed for both EDIL3 and MFGE8 could be due to post-translational modification such as glycosylation (29). An immunoband was still visible in uterine fluid lacking vesicles, indicating that MFGE8 could also be soluble in uterine fluid, unlike EDIL3.

### Immunofluorescence in uterus for key proteins proposed for UF vesicular transport

Immunofluorescence analysis was performed on chicken uterus sections to localize six proteins (ANXA1, ANXA2, ANXA8, CA4, EDIL3, PDCD6IP) proposed to play key roles in vesicular transport within the tissue responsible for eggshell mineralization (Fig. 5). Analyses were performed at five defined time points in the eggshell mineralization process (5 h, 6 h, 7 h, 10 h and 16 h p.o.). Negative controls demonstrated the absence of nonspecific signals in uterus for all antibodies. According to immunofluorescence observations, EDIL3 and ANXA1 protein levels were maximum at 5 h and 6 h p.o., and then decreased from 7 h to 16 h p.o. (Fig. 5). EDIL3 is present in the tubular glands of the lamina propria, whereas ANXA1 was localized in the epithelium. A faint immunostaining was obtained for ANXA2 at 5 h p.o., while the signal was stronger at 6 h, 7 h and 10 h p.o. in the epithelium (Fig. 5), then decreased at 16 h p.o.. ANXA8 was detected in both tubular glands and epithelium regions at each stage of mineralization, with a maximum signal at 10 h p.o. (Fig. 5). A strong CA4 signal was observed in the epithelial cells at 5 h, 6 h and 7 h p.o. while the signal was low at 16 h p.o.. PDCD6IP immunofluorescence revealed that the protein was largely localized in the epithelium throughout the biomineralization process (5 h to 16 h p.o.) (Fig. 5).

**Figure 5.**
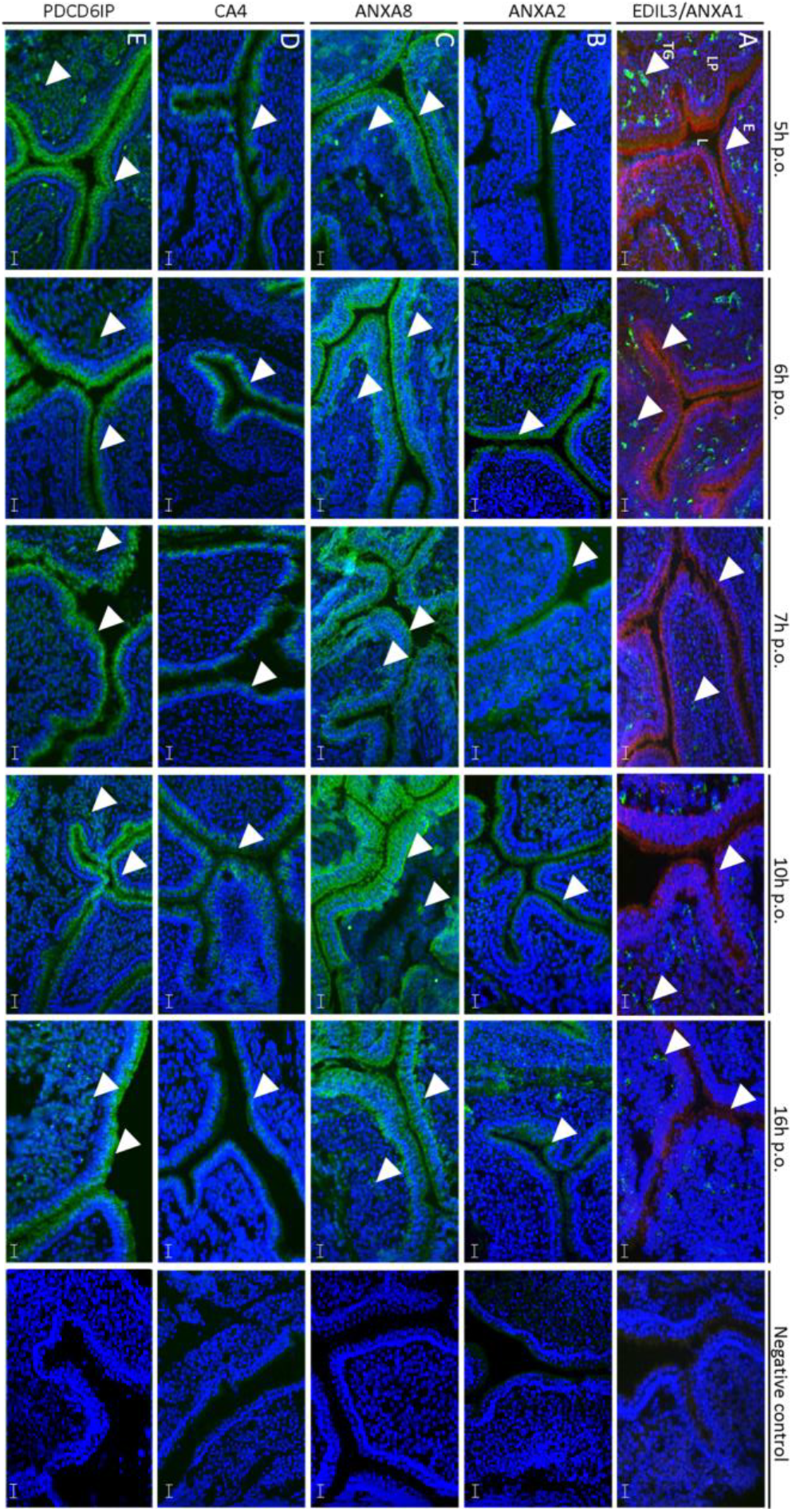
Immunofluorescence of ANXA1, ANXA2, ANXA8, EDIL3, CA4 and PDCD6IP in chicken uterus. The respective rows correspond to (A) co-staining of ANXA1 (red) and EDIL3 (green), (B) staining of ANXA2 (green), (C) staining of ANXA8 (green), (D) staining of CA4 (green) and (E) staining of PDCD6IP (green). ANXA1, ANXA2 and CA4 were localized in the epithelium. ANXA8 and PDCD6IP signals were observed in both epithelium and tubular glands of lamina propria. EDIL3 was solely detected in tubular glands. E: ciliated and glandular epithelium, L: lumen, LP: lamina propria, TG: tubular glands. The primary and secondary antibodies are compiled in Table S5. Bars= 100 μm. White arrowheads indicate positive signal in either tubular glands or epithelium.

## Discussion

Observations of vesicular transport of mineralizing ions during biomineralization have been reported in both invertebrate and vertebrate systems. The most documented vesicular mediated mineralization systems are in vertebrate skeleton and teeth, for which vesicles associated with calcium phosphate have been described (41–43). However, vesicular mediated calcium carbonate mineralization has only been reported in invertebrates. Sea urchin spicules are the best documented example of invertebrate calcite biomineralization mediated by vesicles (5,44). The involvement of vesicles was also proposed for molluscan shell formation (9) and coral exoskeletons (10).

Five to six g of calcium carbonate are deposited within an 18 hours period during chicken egg formation. This requires rapid delivery of large amounts of CaCO_3_, as the laying hen exports 10 % of its total body calcium (2 g) each day (26). Neither component (Ca^2+^ or HCO_3_^−^) is stored in the uterus, but rather are continuously supplied during eggshell formation from the blood plasma *via* transport across the uterine glandular cells. Mechanisms of transepithelial transfer of calcium and bicarbonates have been widely studied and represent the current accepted model for eggshell mineralization (23,26,27). Recently, intestinal paracellular transfer of calcium during eggshell formation was demonstrated (45), raising the possibility of paracellular transfer of calcium in uterus as well; however, no experimental data addressing this possibility is yet available. The vesicular transport of calcium is a third complementary pathway to transfer stabilized ACC to the calcification site.

Shell calcification, which takes place in a saturated milieu (uterine fluid) (46), is a specific event that occurs after ACC accumulation on the eggshell membranes (28). We have proposed a novel model for avian eggshell biomineralization involving EVs budding from uterine epithelial cells, that transit through the uterine fluid (UF) and deliver stabilized ACC to the eggshell membranes and subsequent mineralization front. The advantage of this mechanism is that non-specific precipitation in the fluid cannot occur (29). We report here experimental evidence, which demonstrates the presence of EVs containing ACC in UF, and we define some of the molecular actors involved in this process (Fig. 6). In this study, we observed vesicles from 100 to 500 nm in the uterine cell cytoplasm and their accumulation at the apical membranes. We also observed budding from uterine cells to secrete these vesicles into uterine fluid. The size of vesicles observed in both fluid and uterine cells, and their outward budding, confirmed that EVs similar to shedding microvesicles or ectosomes are present. Secretory activity of multivesicular bodies has been described at the hen utero-vaginal junction, where spermatozoa are preserved in specialized glands (47). Additionally, magnum cells also exhibit vesicle containing egg white proteins (20). Therefore, the distinct oviduct segments seem to use EVs for different functions (i.e. sperm preservation, secretion of egg white proteins and secretion of mineral precursors).

**Figure 6.**
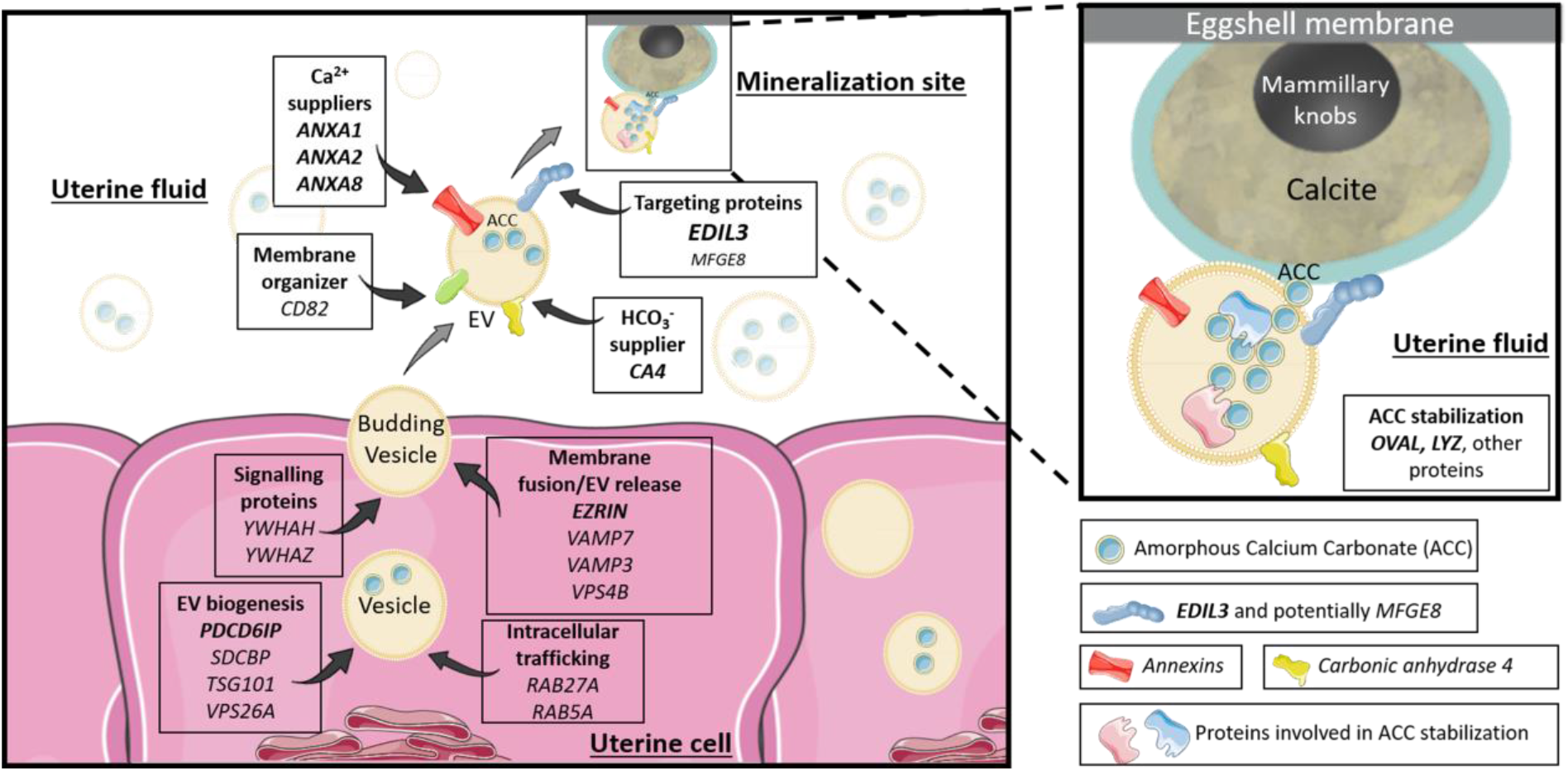
Proposed vesicular function for the different genes and proteins investigated in the study. The extracellular vesicles (EVs) bud by exocytosis from the plasma membrane of the uterine cells. ANXA1, 2, 8 and CA4 supply calcium and bicarbonate ions, respectively (inside uterine cells or in the UF). The EVs transit the uterine fluid (UF) to deliver stabilized ACC (amorphous calcium carbonate) to the mineralization sites (MS). The passage of EV-encapsulated ACC avoids non-specific precipitation in the UF and provides stabilized ACC to the mineralization sites. OVA and LYZ stabilize ACC inside vesicles. EDIL3 (in bold), and to a lesser extent MFGE8, guide the EVs by targeting calcium to the mineralization front. The rest of the proteins are involved in EV biology (biogenesis, intracellular transport and release). The genes in bold represent proteins identified in our proteomic and/or immunofluorescence analyses. Elements were from Servier Medical Art (https://smart.servier.com/), licensed under a Creative Commons Attribution 3.0 Unported License.

This study also gives new important insight on the stabilization and delivery of ACC to the mineralization site where the final structure of eggshell is completed. The avian eggshell is a highly ordered structure, consisting of a porous bioceramic resulting from the sequential deposition of calcium carbonate and organic matrix in the red isthmus/uterus regions of the distal oviduct (17,19,25,48–50). ACC plays a central role in the different phases of shell calcification, as it is a precursor for calcite crystallization during first nucleation events at the mammillary knobs (5 h p.o.) and during eggshell growth (until 22 h p.o.)(28). Eggshell mineralization therefore results from the assembly of aggregated ACC particles and their transformation into calcite crystals to allow a very rapid and controlled process. Additionally, ACC, which is highly unstable under physiological conditions, must be stabilised in this milieu. We have hypothesized that vesicles would allow the transportation of calcium carbonate in amorphous form to the mineralization site (29). In the present study, both EELS and EDS spectroscopy analyzes detected calcium, oxygen and carbon inside EVs. EELS detected carbon K-edge peaks specific for the elemental pattern of calcium carbonate. Electron diffraction on EVs containing calcium carbonate did not detect diffraction circles specific for crystalline polymorphs, confirming the amorphous nature of calcium carbonate. Intracellular vesicles containing ACC were previously observed inside sea urchin epithelial cells (5,51,52). In vertebrates, amorphous calcium phosphate (ACP) has been observed *in vivo* inside intracellular vesicles in embryonic mouse osteoblasts (53). Extracellular vesicles with mineral were also reported for embryonic chick femur (54) and growth plate cartilage of chicken (55,56). As an extension of these observations, we suggest that ACC is packaged inside vesicles when they are formed within the uterine cells. In other biomineralization models, it is known that intracellular calcium is stored in the endoplasmic reticulum (57). However, the subcellular source of ACC-containing EVs detected in our study remains unclear.

The experimental evidence from this study is summarized in a diagram of ACC vesicular transport to the mineralizing site, in which the main identified vesicular actors are incorporated (Fig. 6). We investigated genes involved in ion accumulation / mineral transport by EVs. They included annexins, which can function as calcium channels (58–61), to enable uptake of the calcium ions required for intravesicular ACC formation. Three of them (*Anxa1, -2* and -*8*), exhibited a high expression where eggshell calcification takes place (Ut and RI) (Fig. 7). We also reported in this study that ANXA1, -2 and -8 proteins were also observed in uterine tissue and were found to be overabundant at the initial phase of shell mineralization (5 to 10 h p.o.), compared to later stages. Additionally, ANXA1 (23,62), ANXA2 (23,27,63–65) and ANXA8 (62,65) have already been reported in chicken eggshell proteomic and transcriptomic surveys (62). ANXA1, ANXA2 and ANXA8, exhibit phospholipid-binding domain, and consequently could bind to EV membrane (66). These three annexins constitute, therefore, excellent candidates as participants in vesicular transport, where they could mediate the entry and accumulation of calcium.

**Figure 7.**
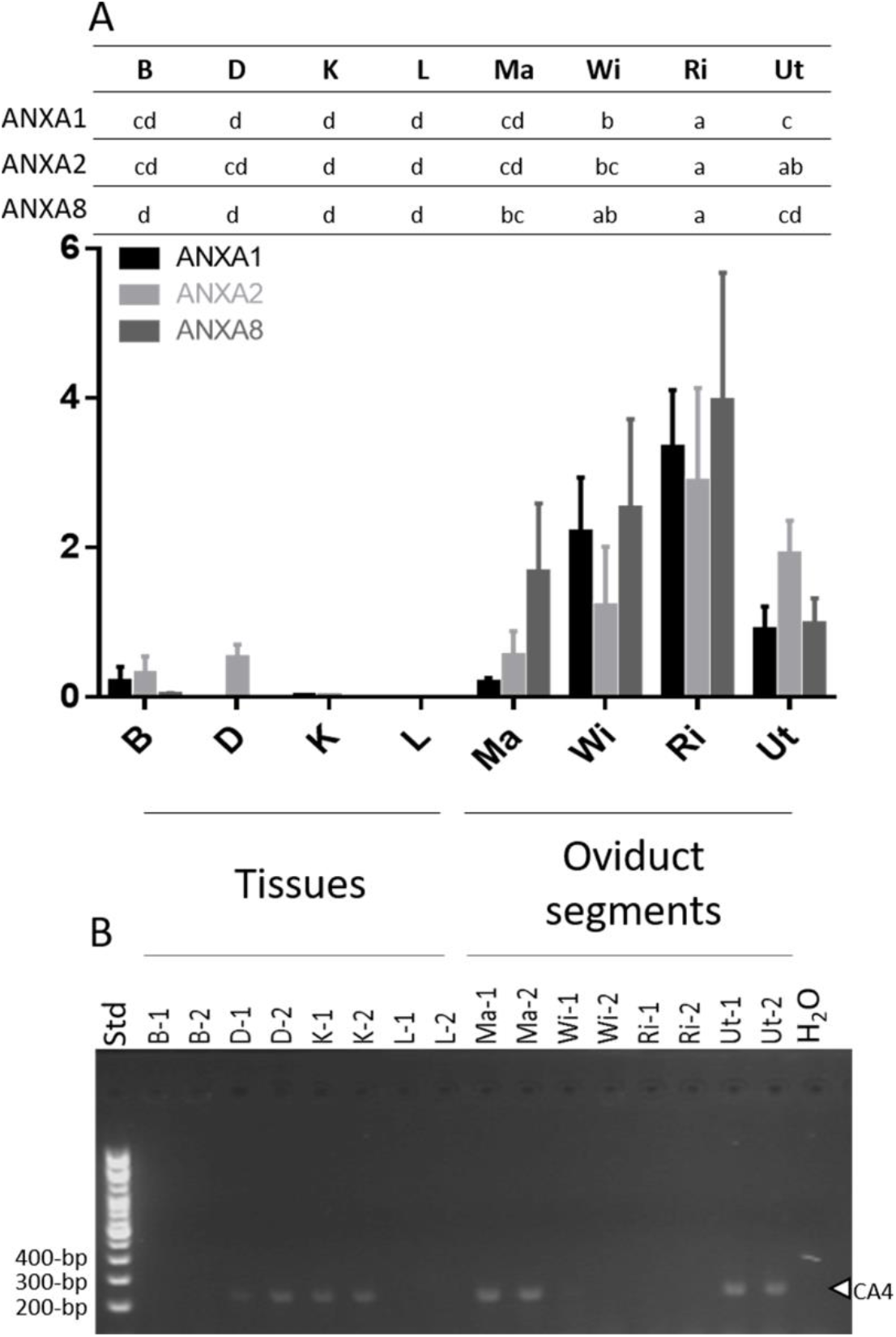
Gene expression of annexins (*ANX*) and carbonic anhydrase 4 (*CA4*). (A) *Anxa1, Anxa2* and *Anxa8* expression. Following reverse transcription, gene expression was quantified using the Biomark microfluidic system (BMK-M-96.96; Fluidigm). Relative quantification was normalized with eight housekeeping genes using GenNorm software. (B) *Ca4* reverse transcription-PCR (100-bp DNA ladder). Gene expression was assessed during the active growth phase of mineralization 10 h p.o. except for B at 18 h p.o.. Tibial bone, B; duodenum, D; kidney, K; Liver, L; Magnum, Ma; White isthmus, WI; Red isthmus, RI; Uterus, Ut; H_2_O, negative control; Std, size (bp) of DNA ladder (100-bp). Letters at the top indicate statistically significant differences of ANXs expression between the tissues (ANOVA-Tukey). The gene accession numbers and primers are compiled in Table S4. GraphPad Prism software was used for diagram representation.

Bicarbonate is a precursor for the calcium carbonate of avian eggshells, and consequently we investigated carbonic anhydrase 2 and 4 (CA2 and CA4) and sodium bicarbonate cotransporter 3 (SLC4A7), as bicarbonate suppliers in vesicular transport, since these have previously detected in uterine transcriptomes during eggshell calcification (23,27). In our study, *Ca2* was not significantly overexpressed in tissues where the shell is mineralized (RI and Ut). We therefore investigated CA4, which was previously identified in chicken eggshell proteomes (62,63,67). In order to confirm its presence in tissues involved in shell mineralization, *Ca4* expression was also measured in the various tissues and organs using end-point RT-PCR (Fig. 7). *Ca4* was expressed in duodenum, kidney and magnum, in addition to Ut where eggshell calcification occurs. The presence of CA4 protein in the uterine epithelial cells was confirmed by immunohistofluorescence, and was overabundant at the onset of shell calcification compared to later stages. Finally, proteomic analysis of purified UF EVs confirmed that CA4 is present in vesicles. CA4 is a glycosylphosphatidylinositol (GPI)-linked protein (68), suggesting that this protein could be anchored to the outside of vesicle membranes and catalyze the reversible hydration of CO_2_ into HCO_3_^−^ at this location. The CA4-catalysed accumulation of high levels of bicarbonate in proximity to the vesicular stores of Ca would promote the formation of intravesicular ACC. SLC4A7 is a sodium bicarbonate cotransporter containing transmembrane domains (https://www.uniprot.org/), which was identified in matrix vesicles released from mineralizing human osteoblast-like cells (35). Its gene was reported in chicken uterine RNA-Seq (https://anr.fr/Projet-ANR-13-BSV6-0007). We also investigated its expression in the oviduct segments but was not significantly overexpressed in RI and Utɞ.

In this study, we quantified the expression of numerous additional vesicle markers within uterine epithelial cells. Among the 20 vesicular genes highly expressed in RI and Ut, seven encode for proteins ranked in the Top 100 of EV protein markers by Vesiclepedia (Table S2). These seven proteins encoded by *Pdcd6ip*, *Hspa8, Anxa2, Ywhaz, Sdcbp, Tsg101* and *Anxa1* are respectively reported to rank 1^st^, 3^rd^, 5^th^, 13^th^, 19^th^, 39^th^ and 43^th^ by Vesiclepedia (40) and were previously reported as constituents of the avian eggshell proteome (Table S2) (62–65,69–78). Moreover, 12 vesicular genes (*Anxa1, Anxa2, Cd82, Edil3, Hspa8, Pdcd6ip, Rab5a, Sdcbp, Tsg101, Vamp7, Ywhah* and *Ywhaz*) encode proteins previously detected in exosomes or matrix vesicles of bone cells, mineralizing cartilage or articular cartilage (Table S2) (32–35,79). The 20 genes reported as highly expressed in the oviduct region responsible for eggshell mineralization encode proteins that have been incorporated into our eggshell vesicular transport model (Fig. 6; Table S2). Some are involved in the general functioning of EVs such as vesicle formation and/or trafficking. Indeed, the genes *Pdcd6ip*, *Sdcbp, Tsg101, and Vps26a* encode proteins involved in the biogenesis of EVs (33,36–39). Proteins encoded by *Rab5a, Rab27a, Vamp7, Vamp3* and *Vps4b* participate in membrane fusion / vesicle release (33,36,37,39). *Yhwah* and *Ywhaz* encode signaling proteins (33), while *Hspa8* and *Cd82* encode chaperone and membrane organizer proteins, respectively (33,37,39). Our proteomics analysis also confirmed the presence of programmed cell death 6-interacting protein (PDCD6IP) and syntenin-1 (*Sdcbp*) in purified vesicles (Table 2 and 3). Syntenin-1 (*Sdcbp*) is a signaling protein and a biogenesis factor described as an extracellular vesicle marker in Vesiclepedia (http://microvesicles.org/extracellular_vesicle_markers). In addition, immunofluorescence for PDCD6IP revealed its presence at each stage and at higher levels during the earliest stage. PDCD6IP is a biogenesis factor for EVs, and is the most frequently reported extracellular vesicles marker (Vesiclepedia). Ezrin is another common EV marker, which is involved in plasma membrane/cytoskeleton cross-linkage during vesicle formation (40,80); it is present in uterine fluid EVs (Table 3). The biogenesis factor, syntenin-1 (*Sdcbp*) and ezrin could be attached to EV membrane *via* their phosphatidylinositol 4, 5 bisphosphate-binding site (Table 3). These results are also supported by previous identification of PDCD6IP, syntenin-1, ezrin in bone and cartilage EVs (32–34).

In our previous study, we underlined the major role of EDIL3 and MFGE8 as proteins predicted to target vesicles containing ACC cargo to the mineralization site (29). EDIL3 and MFGE8 are the 6^th^ and 34^th^ most abundant eggshell proteins, respectively (63). EDIL3 was previously identified in the proteome of cartilage EVs (34). We have previously reported that *Edil3* is overexpressed in the oviduct region where eggshell mineralization takes place (Ut), compared to other oviduct segments or tissues (29). In this study, we identified EDIL3 in the proteomic analysis of purified EVs (Table 2, 3 and Table S3). Using Western blotting, we demonstrated that EDIL3 is only present in the EV fraction of uterine fluid, confirming that this protein is vesicle specific (Fig. 4B). EDIL3 is synthesized by the tubular glands of the uterus (Fig. 5) and could bind to vesicles *via* its phosphatidylserine-binding functionality. In addition, EDIL3 was more abundant during the initiation of mineralization (5-10 h p.o., Fig. 5) (63). Concerning MFGE8, which is paralogous to EDIL3, we observe a high expression of its mRNA in RI compared to other tissues; only liver exhibits a higher expression (29). MFGE8 was not detected in our proteomic analysis of EVs, although Western blotting revealed its presence in UF and to a lesser extend in EVs (Fig. 4C). Altogether, these results strongly suggest that EDIL3 is a key protein in EVs transport at the initiation stage of eggshell biomineralization, while the role of MFGE8 in vesicles transporting ACC remains to be clarified.

We propose two main hypotheses to explain the presence of stabilized ACC in uterine EVs, which are not mutually exclusive. Firstly, accumulation of ACC in the interior of an EV is sufficient to ensure its stabilization, as reported *in vitro* for ACC nanoparticles stabilized in liposomes (81,82). In this case, the membrane coats ACC and shields it from bulk water; reduced water content in the ACC milieu is sufficient to delay crystallization (83,84). Under encapsulated conditions, the initial ACC remains isolated from the aqueous milieu and is observed to dehydrate to the more stable anhydrous ACC phase (82). Moreover, magnesium and phosphate ions are also reported to enhance the stability of ACC and to delay its crystallization (83–85). Physicochemical vesicular conditions could therefore be a key element for ACC stabilization in uterine fluid EVs during eggshell biomineralization.

An alternative hypothesis incorporates additional molecules to thermodynamically stabilize the transient ACC phase (28). Indeed, previous studies have shown that organic matrix components of sea urchin spicules, coral skeleton or mollusk shell can stabilize ACC (86–88). Moreover, ACC was observed using cryo-electron microscopy inside the vesicles of the sea urchin cells involved in spicule mineralization (51,52). In this context, we paid particular attention to lysozyme (LYZ) and ovalbumin (OVA) that were identified in our proteomic study (Table 2 and 3), and were previously reported in chicken eggshell (62,67,69–71,73,89) and shown to modify calcite crystal morphology *in vitro* (90,91). The role of lysozyme in the stabilization of ACC has been explored but remains controversial. Only high concentrations of lysozyme alter the morphology of in *vitro* grown calcite crystals (91). Lysozyme from quail eggshells did not induce the precipitation of ACC under *in vitro* conditions (92). Lysozyme protein was shown to be ineffective in the stabilization of ACC particles (93). However, metastable ACC was obtained *in vitro* in the presence of chicken egg white lysozyme C (94,95). Lysozyme markedly decreased the average diameter of metastable ACC particles and promoted a network of associated particles that incorporated the protein into the precipitate (94). Additionally, lysozyme-ACC particles transform exclusively into the crystalline calcite polymorph (94,95).

OVA, OVALX and OVALY belonging to the serpin family, and have been previously identified in the eggshell matrix (63,90). Their role in calcium carbonate formation and ACC stabilization has been investigated (90,96–99). Calcium binds to ovalbumin and this accumulation creates a nucleation center for mineral formation (99). A model for calcium carbonate mineralization in the presence of ovalbumin has been proposed (97). Calcium ions are bound to the protein by complexation through acidic groups leading to protein rearrangement. The calcium cations are the starting points for the subsequent formation of ACC nuclei, which then undergo a series of phase transitions to the stable crystalline polymorphs (98). The ability of OVA to stabilize unstable calcium carbonate phases was further confirmed through *in vitro* experiments (99,100). OVA is one of the major proteins involved in shell formation and its presence in vesicles will contribute to the formation of metastable ACC (Table 3, Fig. 6).

We also identified about 20 additional proteins in UF EVs by proteomic analysis (Table 2 and Table S3). Ovotransferrin (OVOT) is known to modify calcite crystal morphology at low concentration (101). Alpha-2-macroglobulin (A2M), another protein identified in our study (Table 2 and Table S3), is an egg yolk protein able to interact with low-density lipoprotein, which exhibits binding calcium properties and consequently could interact with ACC (https://www.uniprot.org/) (20,102). Ovomucin (MUC), ovomacroglobulin (ovostatin, OVST) and ovoglobulin G2 (TENP or OVOG2) are egg white proteins, whereas vitelline membrane outer layer protein (VMO1) is a protein enriched in the vitelline membrane (Table 2 and Table S3) (103). They were previously identified in eggshell (62,63), but their role related to the calcification process is unknown. Clusterin (CLU) is a chaperone protein previously identified in eggshell and proposed to prevent the premature aggregation and precipitation of eggshell matrix proteins during calcification (104). Aminopeptidase N (AMPN) and Loc 771972 (OCX25) are a protease and protease inhibitor, respectively (17,105), which were also detected in vesicles (Table 2 and Table S3). Protease and protease inhibitors could potentially control the calcification process, either by degrading proteins or by modifying the processing of protein maturation (63). Prominin-1-A (PROM1), a protein involved in lipid metabolism was also present in vesicles but its role is not yet defined. Finally, nine EV proteins are partial immunoglobulins, with unknown functions in vesicles or in shell mineralization.

We consequently propose a comprehensive model for calcium and carbonate transport to the mineralization site during eggshell formation (Fig. 8). Calcium and carbon dioxide originate from blood. Blood CO_2_ passively diffuses into uterine cells (106), where it will be hydrated by CA2. Alternatively, bicarbonate can be actively transferred into uterine cells using the Na^+^/HCO_3_^−^ co-transporters SLC4A4-A5-A10 (26). Bicarbonates are actively extruded from cells by the HCO_3_^−^/Cl^−^ exchanger SLC26A9 (26). Additionally, bicarbonate ions will be produced in uterine fluid by hydration of CO_2_ by CA4, which has its active site in the extracellular space (107). The transcellular pathway to secrete calcium and bicarbonate ions into the fluid has been previously described (23,27). TRPVs (transient receptor potential cation channels) and/or otopetrin 2 (OTOP2) and ATPase secretory pathway Ca^2+^ transporting 2 (ATP2C2) would allow the entry of plasma calcium, and it is buffered / transferred intracellularly by calbindin-1 (other Ca^2+^ pumps associated with the endoplasmic reticulum could be also involved in this transfer); additionally, the Ca^2+^/Na^2+^ exchangers SLC8A1-3 and the Ca^2+^ pumps ATP2B1-B2 are involved in the apical extrusion of calcium into the uterine fluid (26,108). A paracellular Ca^2+^ uptake pathway was recently described in intestine that acts to replenish calcium stores from dietary sources during eggshell biomineralization. The paracellular pathway involves claudins (CLDN), occludins (OCN), junctional adhesion molecules (JAM) and tight junction proteins (TJP) (45). Our uterine RNA-Seq analysis reveals elevated expression of several genes of this paracellular pathway (*Tjp1*, *Cldn1*, *Cldn10*, *Ocln*, *Jam2*) (https://anr.fr/Projet-ANR-13-BSV6-0007). Moreover, expression of *Cldn10* has also been detected in chicken uterus (108,109). Although the involvement of the paracellular pathway in eggshell calcification is controversial, it has been incorporated in our comprehensive model of ionic pathways during shell biomineralization (Fig. 8). The ionic calcium concentration in uterine fluid ranges from 6 to 10 mmol/l depending of the stage of calcification (46), which is higher than blood ionic calcium levels (1-2 mmol/l); consequently the concentration gradient is not in favor of calcium movement towards the uterine fluid (26). However, Bar (2009) suggested that the electrical potential difference could invert this gradient, allowing paracellular transfer of calcium into the uterine fluid (110). Moreover, the concentration of potassium in uterine fluid (10 to 65 mM dependent on stage of mineralization) is higher than in the blood plasma (4 mM) (23). Consequently, the paracellular pathway could participate in potassium transfer to maintain ionic homeostasis. Finally, we have added our evidence related to vesicular ACC transport within uterine fluid. Based on the available literature, we propose that ACC is taken up by vesicles that form inside uterine epithelial cells (1 in Fig. 8). Intracellular vesicles containing stabilized ACC are then secreted into the uterine fluid to be targeted to mineralization sites. Alternatively, vesicles could be secreted without ACC; if so, ACC would accumulate in vesicles within the uterine fluid (2 in Fig. 8). Further experiments will be necessary to distinguish between these possibilities. In both cases, we propose that Annexins present on vesicle membranes promote calcium entry, whereas CA4 catalyzes the hydration of CO_2_ into bicarbonate ions. ACC inside EVs are delivered to the mineralization site, with EDIL3, and possibly MFGE8, as guidance molecules for this targeting (29). Additional calcium / bicarbonate ions provided by transcellular, and possibly paracellular, pathways would allow the epithaxial growth of calcite at the mineralization front during initial and subsequent stages of shell mineralization.

**Figure 8.**
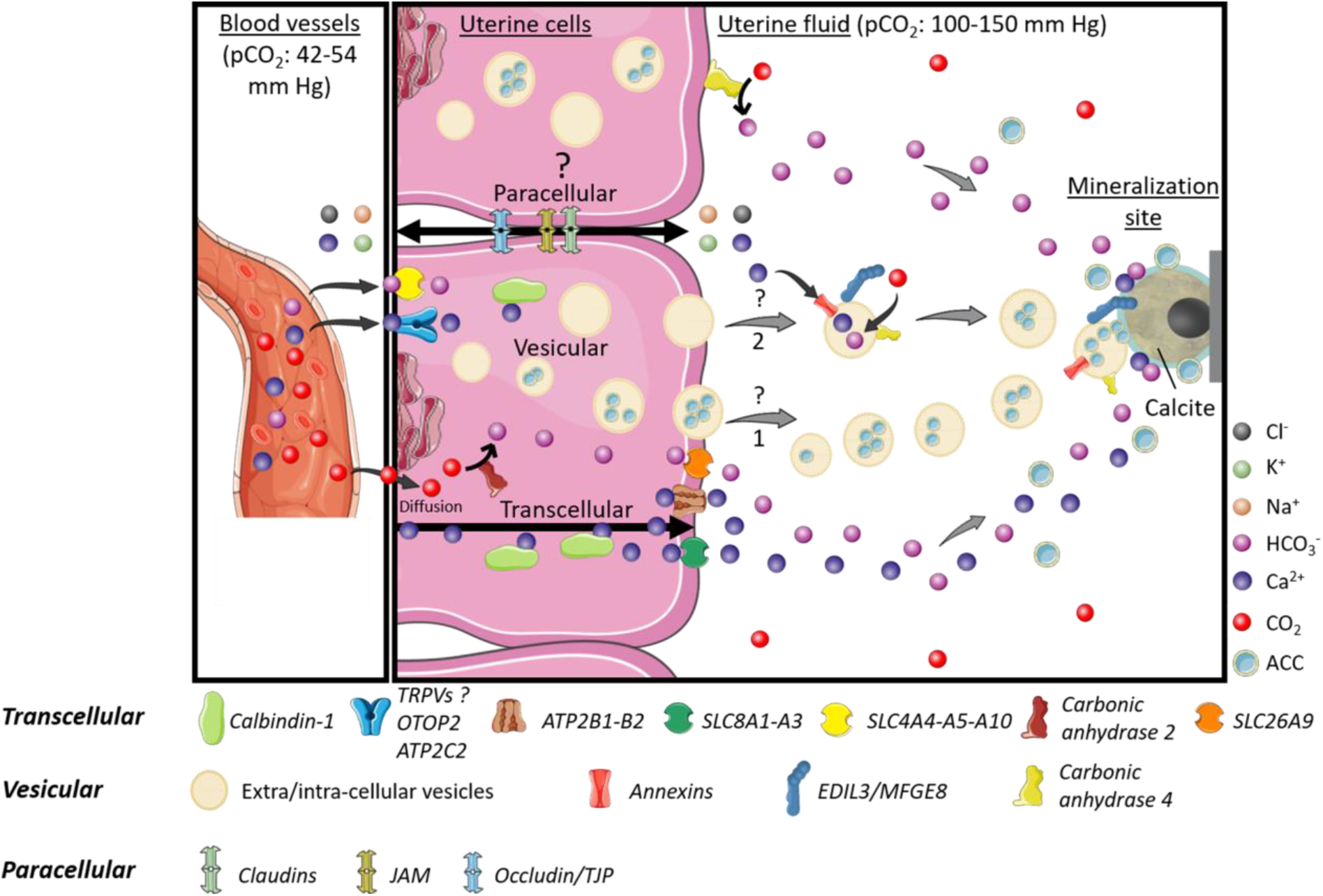
Model of calcium and carbonate transport to the uterine fluid during eggshell calcification. The three potential pathways for ion transfer through uterine cells are transcellular, vesicular and paracellular mechanisms. They could function asynchronously or in an integrated fashion. The major protein players in each pathway are indicated at the bottom of the figure. In the vesicular pathway, vesicles could accumulate ACC either inside the uterine cells (1) or in the uterine fluid (2). Elements were from Servier Medical Art (https://smart.servier.com/), licensed under a Creative Commons Attribution 3.0 Unported License.

In a previous study, we proposed a vesicular transport model that could deliver ACC for deposition during eggshell biomineralization (29). In contrast to the eggshell, which is found only in birds and some reptiles, all vertebrates possess otoconia or otoliths as calcium carbonate biomineralized structures necessary for balance and sensing linear acceleration (111,112). A previous study highlighted similarities between proteins involved in eggshell and otoconia formation (67); however, the potential role of vesicular mediated transport to produce them remains unknown. Our results on vesicular transport of ACC could provide insight into physiological mineralization of otoconia. In the present study, we provide evidence for the first time in vertebrates of a vesicular transport pathway for delivery of stabilized amorphous calcium carbonate. We used multiple approaches to confirm predictions based on this model. TEM coupled to elemental analysis demonstrated several phases of EV formation at uterine cells, as well as the presence of EVs containing ACC in the UF. The major actors involved in this vesicular transport were organized into a comprehensive schema (Fig. 6), which incorporates EVs and constituent proteins in the process of shell mineralization. EDIL3 and possibly MFGE8 are targeting proteins that are proposed to address vesicles for delivery of ACC to the mineralization site. Calcium accumulation by vesicles is promoted by annexins 1, 2 and 8, whereas carbonic anhydrase 4 catalyzes bicarbonate formation from CO_2_. Moreover, our study identified and characterized several additional proteins that could contribute to the stabilization of ACC inside vesicles, under favorable physical and chemical conditions. Finally, we propose an updated and coherent model for the transfer of ions to ensure avian eggshell biomineralization (Fig. 8). This model blends together transcellular, paracellular and vesicular pathways to address the supply of minerals necessary to build this composite biomaterial. Further work will be necessary to define the relative contribution of each pathway.

## Experimental procedures

### Ethical statement and housing

All animal-handling protocols were carried out in accordance with the European Communities Council Directives concerning the practice for the care and use of animals for Scientific Purposes and the French Ministry on Animal experimentation, and were performed under the supervision of authorized scientists (authorization #7323, delivered by the French ministry of Education and Research). Birds were kept in the experimental unit PEAT 1295 of INRA, which has permission to rear birds and to perform euthanasia of experimental animals (decree N° B37-175-1 of August 28^th^ 2012 delivered by the “Préfécture d’Indre et Loire” following inspection by the Direction of Veterinary Services). Our experimental protocol was approved by the ethical committee “Comité d’éthique de Val de Loire, officially registered under number 19 of the French National Ethics Committee for Animal Experimentation” under agreement number #16099-015902.

Mature brown egg-laying hens (ISA-Hendrix, 40 and 90 weeks old at sampling) were kept in individual cages furnished with automatic equipment for recording of the oviposition (egg-laying) times. Animals were fed layer mash *ad libitum* and exposed to a 14 h light / 10 h darkness cycle. Following collection and removal of egg contents, the eggshells were thoroughly washed with water, air dried and stored at −20°C.

### Tissue and uterine fluid collection

Thirty-two brown laying hens (ISA-Hendrix, 26 and 6 at 40 and 90 weeks old, respectively) were sacrificed by lethal intravenous injection of Dolethal® (Vetoquinol, Magny-vernois, France) at either the initial phase of eggshell mineralization (5-7 h p.o.), the onset of the active mineralization phase (10 h p.o.) or during the linear growth phase of rapid mineralization (16-18 h p.o.).

For RT-PCR and Western blotting analyses, various tissues (Duodenum, D; Kidney, K; Liver, L) and oviduct regions (Magnum, Ma; White isthmus, WI; Red isthmus, RI and Uterus, Ut) were collected from 40 week old hens sacrificed at 10 h p.o., whereas mid-shaft tibial bones (B) were collected at 18 h p.o from 90 week old hens. Tissues were directly immersed in liquid nitrogen and then stored at −80 °C in cryo-tubes until RNA or protein extraction.

Small pieces of uterine wall (0.5 cm^2^) were also sampled at 16 h p.o. from 40 week old hens for electron microscopy, and were fixed by incubation at room temperature for 24 h in 4% paraformaldehyde, 1% glutaraldehyde in 0.1 M phosphate buffer (pH 7.3). Fixed tissues were stored at 4°C until preparation of sections.

For immunofluorescence analysis, small pieces (2 cm & 0.4 cm) of the uterus at each stage (5 h, 6 h, 7 h, 10 h and 16 h p.o.), from 25 hens at 40 week old, were embedded in O.C.T. (Optimal cutting temperature) compound mounting medium for cryosectioning (VWR, Radnor, USA) using molds and direct immersion in isopentane cooled by liquid nitrogen. The blocks of O.C.T. containing uterus pieces were stored at −80 °C until cryosectioning.

UF was collected as previously described (22). Briefly, egg expulsion was induced by injection of 50 μg prostaglandin-F2α in the alar vein of 40 week old hens, either at 9 h or 16 h p.o. Following egg expulsion, UF was collected directly by gravity in a plastic tube placed at the entrance of the everted vagina. A portion of the fluid was promptly pipetted and deposited in liquid nitrogen to obtain 50 μL beads of uterine fluid for storage at −80 °C until use for electron microcopy. The remaining UF was diluted in phosphate buffered saline (PBS, 4 v: 1 v) in cryotubes maintained on ice and immediately used for extracellular vesicle isolation.

### Gene expression

Total RNA was extracted from tissues harvested at 10 h p.o. (except bone, which was extracted at 18 h p.o.), and treated as previously described (29). Briefly, the concentration of each RNA sample was measured at 260 nm with NanoDrop™ and their integrity was assessed on 1.5 % agarose gels. Total RNA samples (1 μg) were reverse-transcribed using RNase H-MMLV reverse transcriptase (Superscript II, Invitrogen, Carlsbad, California) and Oligo (dT)™ primers (Invitrogen, Carlsbad, California). Primers for the *Edil3*, *Mfge8*, *Ca4*, the 31 vesicular genes and the eight housekeeping genes (*B2m, Eif3i, Gapdh, Gusb, Stag2, Tbp, Sdha*, and *Ppia*) were designed from *Gallus gallus* gene sequences (Table S4), using Primer-BLAST on National Center for Biotechnology Information (NCBI), and were synthesized (Eurogentec, Lièges, Belgium). Primer efficiencies were evaluated by RT-qPCR using LightCycler^®^ 480 SYBR Green I Master and LightCycler^®^ 480 instrument II (Roche, Bâle, Switzerland). Gene expression was quantified using the Biomark microfluidic system, in which every sample-gene combination is quantified using a 96.96 Dynamic Array™ IFCs (BMK-M-96.96, Fluidigm^®^, San Francisco, California) at the GeT-GenoToul platform (Toulouse, France). Samples were pre-amplified according to the manufacturer’s specifications. PCR was then performed using the following thermal protocols: Thermal Mix (50 °C, 2 min; 70 °C, 30 min; 25 °C, 10 min), Hot Start (50 °C, 2 min; 95 °C, 10 min), 35 PCR cycles (95 °C, 15 s; 60 °C, 60 s), and Melting Analysis (60 °C, 30 s; 95 °C, 1 °C/3 s). Real time quantitative PCR results were analyzed using Fluidigm Real-time PCR analysis software v.4.1.3 (https://www.fluidigm.com/software). Six biological replicates and two technical replicates were carried out for each tissue. GenNorm software (https://genorm.cmgg.be/) was used for validation of housekeeping gene stability. The normalized quantities were calculated using the following formula: (gene efficiency ^(ctcalibrator-ctsample)^)/geometric average quantity of the eight housekeeping genes. Normalized quantities were compared between various measured tissues and oviduct segments using one-way ANOVA followed by Tukey pairwise test analysis on Minitab® 18 software (http://www.minitab.com/fr-fr/). A p-value < 0.05 was selected as the threshold for significance between different groups. The heatmap of the gene expression levels (Row-Z score) in different tissues for *Edil3*, *Mfge8* and the 31 vesicular genes was computed using the heatmap.2 function from gplots packages of Rstudio software 1.1.456 (https://www.rstudio.com/). Row and column clustering were performed using pearson-ward.D2 methods and spearman-Ward.D2 methods, respectively.

The *Ca4* mRNA was amplified by end-point PCR in each oviduct segment and tissues using DreamTaq PCR Master Mix 2X (ThermoFisher Scientific, Waltham, USA) and Mastercycler gradient (Eppendorf, Hambourg, Germany). After 3 min of denaturation, 35 cycles were used (95 °C, 30 s; 60 °C, 30 s; 72 °C, 30 s) to amplify PCR products, followed by a final elongation for 15 min. PCR products were then mixed with Trackit cyan/orange Loading buffer 6 X (ThermoFisher Scientific, Waltham, Massachusetts), loaded on a 2.5 % agarose gel and migrated for 30 min at 100 V using the Mupid-One electrophoresis system (Dominique Dutscher, Issy-les-mouligneaux, France). The Trackit 100-bp DNA ladder was used (ThermoFisher Scientific, Waltham, Massachusetts) to determine the size of PCR products. Imaging was performed using the Syngene GeneGenius Gel Light Imaging System (Syngene, Cambridge, UK).

### Extracellular vesicle isolation from uterine fluid and extraction of soluble eggshell matrix proteins

EVs were isolated from uterine fluid harvested at 9 h and 16 h p.o., using previously described methodology (113). Briefly, UF diluted in PBS was centrifuged at 100 & g for 15 min and then at 12, 000 & g for 15 min to remove cell debris. Two successive ultracentrifugation steps at 100, 000 & g (Beckman Coulter L8-70M, Beckman Coulter, Brea, California) were then performed during 90 min to pellet the EVs. Aliquots of the supernatants corresponding to UF without EVs were sampled and stored at −20 °C. The pellet was suspended in 50 μL of PBS and stored at −20° C until transmission electron microscopy (TEM), nanoLC-MS/MS and Western blot analysis.

Soluble eggshell proteins (SEP) from normally laid eggs were extracted as previously detailed (101). Briefly, eggshell pieces with eggshell membranes were immersed in 154 mM NaCl containing protease inhibitors (2.5 mM benzamidine-HCl, 50 mM ε-amino-n-caproic acid, 0.5 mM N-ethymaleimide, and 1 mM phenylmethylsulfonyl fluoride), and then ground into a fine powder. Eggshell powders were fully demineralized by immersion in 20 % acetic acid, and the resulting suspensions were dialyzed (cut off 3,500 Da; dialysis membrane Spectra/Por™, ThermoFisher Scientific, Waltham, Massachusetts) against demineralized water for 24 h at 4 °C and lyophilized. Powdered samples were incubated overnight at 4 °C in 4 M guanidine-HCl, 5 mM benzamidine-HCl, 0.1 M ε-amino-n-caproic acid, 10 mM EDTA, 50 mM sodium acetate and 1 mM phenylmethylsulfonyl fluoride, and then dialyzed (cut off 3, 500 Da) against 0.5 M sodium acetate pH 7.4 for 24 h at 4 °C, followed by centrifugation at 2, 000 & g for 10 min at 4 °C. The resulting SEP (supernatants) were stored at −20 °C.

### Electrophoresis and Western blot analyses

Protein concentrations were determined using the BioRad DC Protein Assay kit II (BioRad, Marnes-la Coquette, France) in accordance with manufacturer’s instructions and using bovine serum albumin (BSA; Sigma-Aldrich, Saint-Quentin Fallavier, France) as standard. Fifteen micrograms of each sample were diluted into Laemmli sample buffer (5v:1v) and boiled for 5 min. Protein samples were then separated on 12 % polyacrylamide gels (Mini-Protean II electrophoresis cell, BioRad, Marnes-la-Coquette, France) and transferred to 0.2 μm nitrocellulose blotting membrane (GE Healthcare, Little Chalfont, UK) for Western blot analysis. Briefly, membranes were washed 5 min in Tris buffered saline (TBS; 50 mM Tris-HCl, 150 mM NaCl, pH 7.4), blocked with the Odyssey® blocking buffer (LI-COR, Bad Homburg, Germany) in TBS (1v:1v), and then incubated for 3 h in the blocking solution (Odyssey® blocking buffer 1v: TBS 1v) containing 0.1% Tween-20 and anti-EDIL3 (1: 1, 000) or anti-MFGE8 antibodies (1: 1, 000) (Table S5). Membranes were sequentially washed for 5 min in TBS, 0.1 % Tween-20, and then incubated for 1 h in the blocking solution containing 0.1 % Tween-20 and AlexaFluor® 680 goat anti-rabbit (H+L) secondary antibody (1: 20, 000; ThermoFisher Scientific, Waltham, Massachusetts) (Table S5). Finally, membranes were washed three times in TBS containing 0.1 % Tween-20 and twice in TBS. Immuno-reactive bands were revealed using the Odyssey® imaging system (LI-COR, Bad Homburg, Germany) at 700 nm.

### Immunofluorescence of uterine tissue

Serial sections (10 μm) were prepared from uterus embedded in O.C.T. (harvested at 5 h, 6 h, 7 h, 10 h and 16 h p.o.) using the CM3050 S cryostat (Leica, Wetzlar, Germany). Sections (stored at −80 °C until use) were thawed for 30 min at room temperature and rehydrated during 5 min in 1X PBS before immunolabeling. The blocking step was performed with 1X PBS, 3 % BSA for 20 min at room temperature. Sections were then incubated in 1X PBS, 3 % BSA with diluted primary antibodies (anti-ANXA1, anti-ANXA2, anti-ANXA8, anti-CA4, anti-EDIL3 or anti-PDCD6IP, Table S5). Sections were washed 3 times at room temperature for 5 min in 1X PBS. Negative controls were prepared without primary antibody. Sections were washed 3 times for 5 min at room temperature in 1X PBS. Then, according to the primary antibody, sections were incubated with either goat anti-rabbit secondary antibody (IgG, H+L), AlexaFluor® 488 (1: 1, 000; ThermoFisher Scientific, Waltham, Massachusetts) or goat anti-mouse IgG1 cross-adsorbed secondary antibody, AlexaFluor® 555 (1: 1, 000; ThermoFisher Scientific, Waltham, Massachusetts) (Table S5) in 1X PBS, 1 % BSA for 1 h at room temperature. All immunolabeling was performed on duplicate sections. Sections were washed three times for 5 min at room temperature and then mounted using Fluoroshield™ with DAPI histology mounting medium (Sigma-Aldrich, Saint-Quentin Fallavier, France). Sections were observed with Axioplan II Fluorescence Microscope (Zeiss, Oberkochen, Germany) and images were acquired using an INFINITY microscope camera (Lumenera, Nepean, Canada).

### Proteomics analysis of uterine fluid extracellular vesicles

Sixty micrograms of purified EVs resuspended in PBS were diluted into 2 % SDS Laemmli sample buffer (5v:1v) and boiled for 5 min. EV proteins were electrophoresed on a 12.5 % polyacrylamide gel (50V, 30 min) to obtain a single band (Mini-Protean II electrophoresis cell, BioRad, Marnes-la-Coquette, France). The gel was stained with Coommassie Blue and the single band of proteins was excised and subjected to in-gel digestion with bovine trypsin (Roche Diagnostics GmbH, Mannheim, Germany) as previously described by Marie *et al.* for analysis by nanoscale liquid chromatography–tandem mass spectrometry (nanoLC–MS/MS) (76). MS/MS ion searches were performed using Mascot search engine version 2.6 (Matrix Science, London, UK) *via* Proteome Discoverer 2.1 software (ThermoFisher Scientific, Bremen, Germany) against the NCBIprot_chordata database (July 2018). The search parameters included trypsin as a protease with two allowed missed cleavages, carbamidomethylcysteine, methionine oxidation and acetylation of protein N-termini as variable modifications. The tolerance of the ions was set to 5 ppm for parent and 0.8 Da for fragment ion matches. Mascot results obtained from the target and decoy databases searches were subjected to Scaffold software (v 4.8.8, Proteome Software, Portland, USA) using the protein cluster analysis option (assemblage of proteins into clusters based on shared peptide evidence). Peptide and protein identifications were accepted if they could be established at greater than 95.0 % probability as specified by the Peptide Prophet algorithm and by the Protein Prophet algorithm, respectively. Protein identifications were accepted if they contained at least two identified peptides.

### Transmission electron microscopy and electron energy loss spectroscopy analyses of uterine fluid

Frozen beads of UF harvested at 16 h p.o were deposited on a Ni electron microscope grid and dried for 1 min. Excess UF was then absorbed with filter paper and the grid was immediately observed with the TEM-ZEISS LIBRA 120 instrument (ZEISS, Oberkochen, Germany). EVs were localized under the TEM and electron energy loss spectroscopy (EELS) spectra from 270 to 410 eV and from 460 to 600 eV were acquired (Table S1). Mapping of elemental calcium (345-355 eV) and carbon (280-290 eV) were performed to determine the presence and organization of CaCO_3_ and organic phase carbon in EVs.

### Transmission electron microscopy and energy-dispersive X-ray spectroscopy analyses of uterus and uterine fluid

Fixed pieces of uterus were washed in PBS and post-fixed by incubation with 2 % osmium tetroxide for 1 h. Samples were then fully dehydrated in a graded series of ethanol solutions and embedded in Epon resin, which was polymerized by heating from 37 °C to 60 °C. Ultra-thin sections (50–70 nm thick) of these blocks were obtained with a LEICA Ultracut UCT ultramicrotome (Leica, Wetzlar, Germany). Sections were stained with 5 % uranyl acetate, 5 % lead citrate and observations were made with a TEM-1400 Plus electron microscope (JEOL, Tokyo, Japan).

UF harvested at 16 h p.o. and stored as frozen beads, was deposited on a Ni electron microscope grid (C coated) and then dried for 1 min. Excess UF was then absorbed with filter paper, stained with uranyl acetate 2 % and the grid was immediately observed with a TEM-1400 Plus electron microscope (JEOL, Tokyo, Japan). EVs were observed under the TEM (BF/DF detector) and energy-dispersive X-ray spectroscopy (EDS; Oxford 65 mm^2^ detector) analysis was performed on EVs in order to detect elemental calcium, carbon and oxygen (AZtec software). Selected area electron diffraction (SAED) analysis was also carried out to determine the crystalline phase of minerals observed in EVs.

### Data Availability

All data are contained within the manuscript.

## Acknowledgements

The authors are grateful to the experimental unit (PEAT, INRAE, 2018. Poultry Experimental Facility, doi: 10.15454/1.5572326250887292E12) for the care of birds, to the GeT-GenoToul platform Toulouse for performing real time quantitative PCR with the Biomark Fluidigm system and to the Centro de Instrumentación Científica de la Universidad de Granada for electron microscopy and elemental analysis. The authors acknowledge Pierre-Ivan Raynal and Sonia Georgeault from the IBiSA electron microscopy platform of the “Université François Rabelais” of Tours for their assistance.The authors also acknowledge Nadine Gérard, Cindy Riou and Agostinho Alcantara-Neto from UMR “Physiologie de la Reproduction et des Comportements” (INRAE Centre Val de Loire) for their help in extracellular vesicle isolation. The authors wish to thank the Université François Rabelais de Tours and the Région Centre Val de Loire for financial support of Lilian Stapane’s doctoral studies. MTH acknowledges funding from NSERC (RGPIN-2016-04410) and is grateful to LE STUDIUM for support during the preparation of this manuscript. He is a LE STUDIUM Research Fellow, Loire Valley Institute for Advanced Studies, Orleans-Tours, and BOA, INRAE, Centre Val de Loire, Nouzilly, France.

## Author contributions

LS was involved in designing and planning of the study. He performed the experiments and analyses, interpreted data and statistical analyses and wrote the original draft of the paper. NLR was involved in the conceptualization of the study, the experimental design and the interpretation of data and contributed to the writing of the paper. JE, ABRN, VL, LCS carried out a part of experiments and interpreted data. MTH was involved in the conceptualization and the writing of the paper. JG conceived the strategy, designed and experiments, interpreted data and statistical analyses and was involved in the writing of the paper. He acquired funding and supervised the study. All authors have read and approved the final manuscript.

## Supporting Information

**Table S1.** Assignments of electron energy loss spectroscopy (EELS) peaks. Approximate energy loss (eV) were deduced for each peak from carbon K-edge, calcium L2,3 edge and oxygen K-edge in the TEM-EELS analysis.

| Element | Energy loss (eV) | Peak assignment |
| --- | --- | --- |
| carbon K-edge | 285.5 | C=C |
|  | 290.3 | carbon-oxygen bonds |
|  | 295.5 | carbon-oxygen bonds |
|  | 301.5 | carbon-oxygen bonds |
| calcium L2,3-edge | 349.3 | Ca L<sub>3</sub> edge |
|  | 352.6 | Ca L<sub>2</sub> edge |
| oxygen K-edge | 534 | C=O bonds |
|  | 540 | CO<sub>3</sub><sup>2-</sup> group |
|  | 545.5 | C=O |

**Table S2.** (**.xlsx file**) Summary of the state of knowledge for different genes and proteins investigated in our study. The file contains genes and proteins potentially involved in vesicular transport for eggshell biomineralization. EV: extracellular vesicle, UF: uterine fluid. Vesiclepedia database (www.microvesicles.org/) classifies the proteins that are frequently identified in EVs. Protein localization in EVs was predicted using molecular function of proteins on Uniprot (https://www.uniprot.org/). References: (24,27,32-35,41-43,58-65,67,69-79,114-120), * https://anr.fr/Projet-ANR-13-BSV6-0007.

**Table S3.** (**.xlsx file**) Proteins identified by proteomics analysis of uterine fluid extracellular vesicles. List of the twenty-nine non-redundant proteins identified by nanoLC-MS/MS analysis on extracellular vesicle (EVs) fraction of the uterine fluid (9 h p.o.).

**Table S4.**
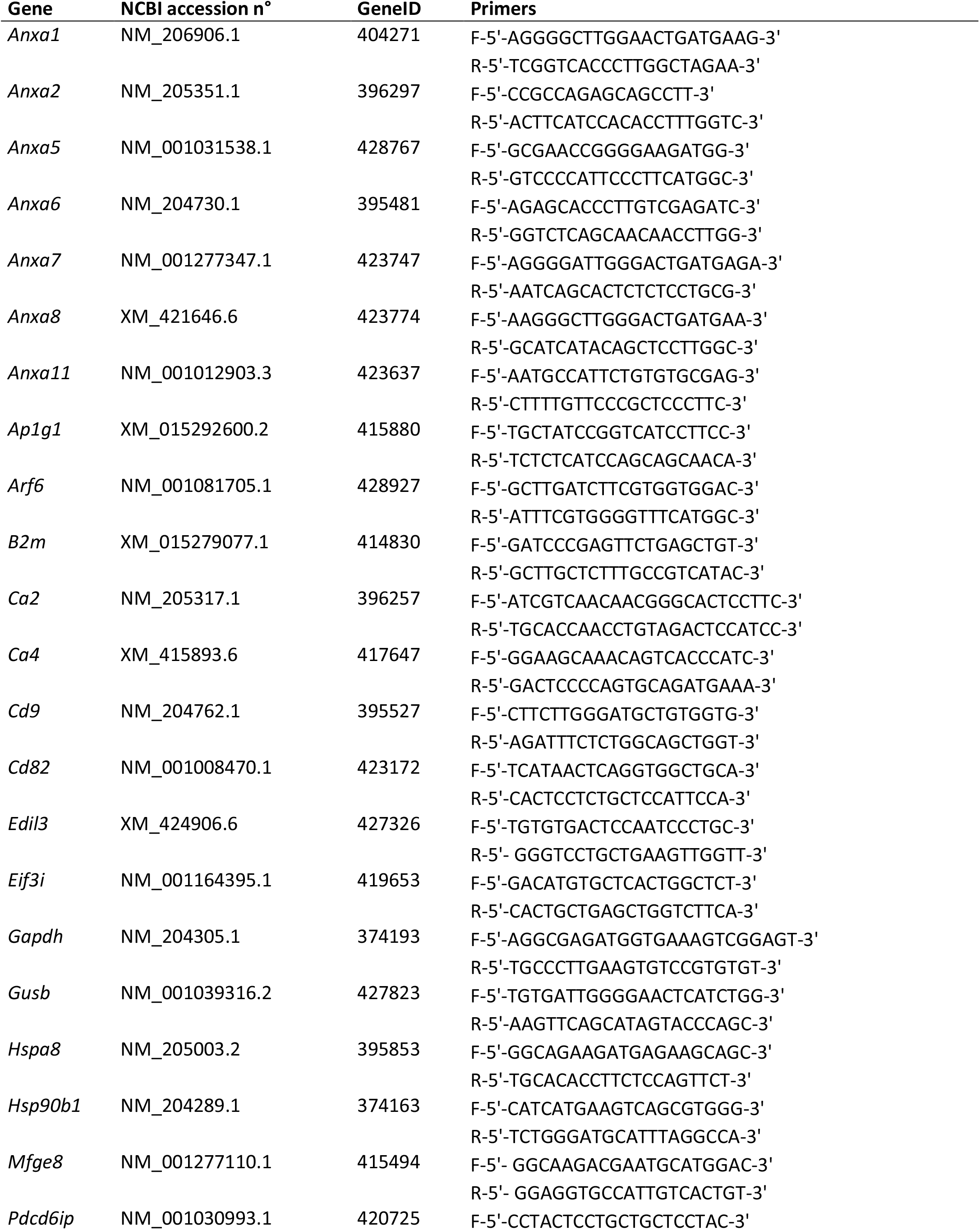
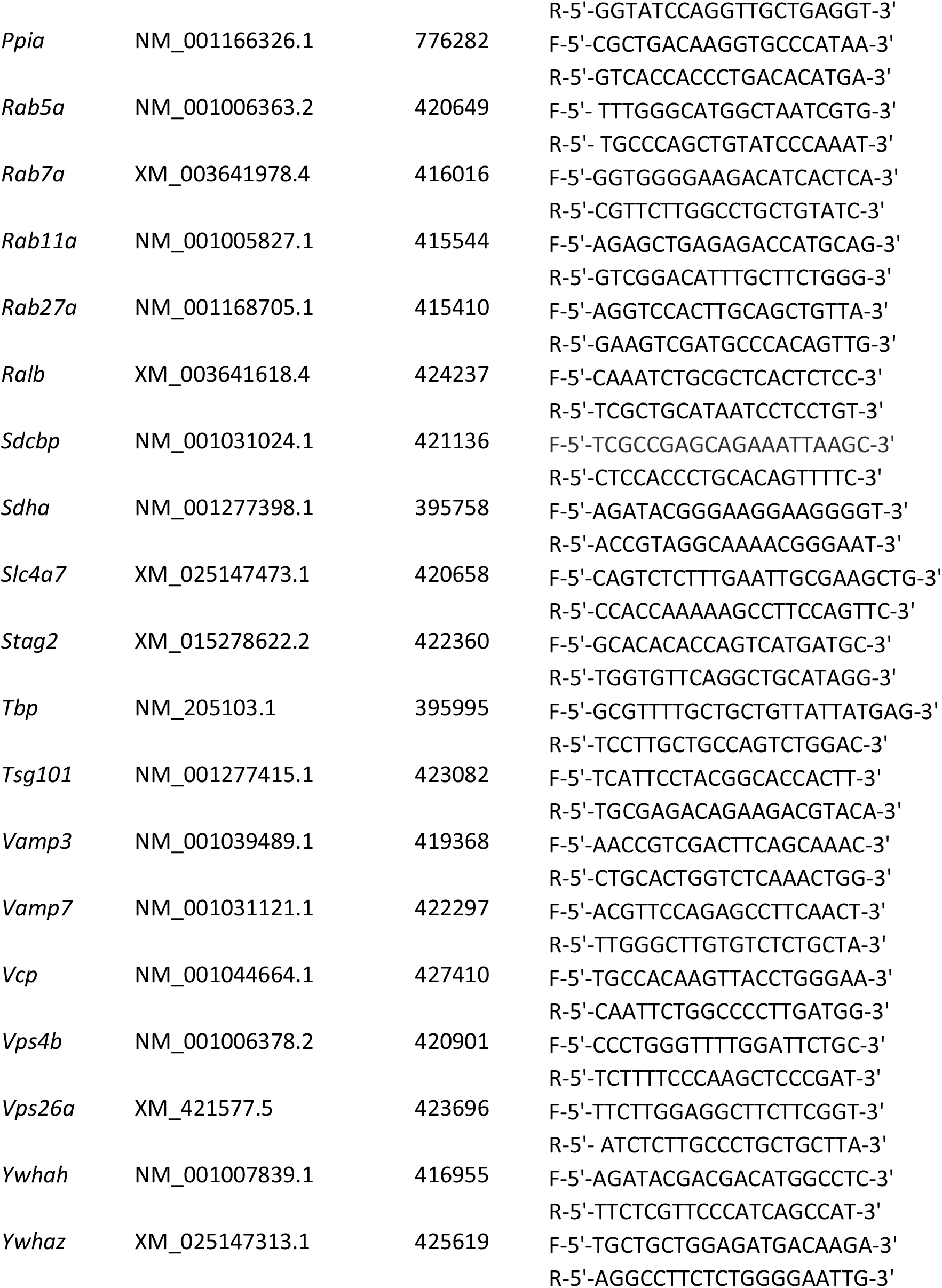
Primers used for the gene expression study. Accessions correspond to GenBank accession numbers.

**Table S5.** Antibodies used in the study. Optimal dilution of antibodies for immunofluorescence (IF) and Western blotting (WB) are indicated.

| Antibodies | Species/type | Dilution<br>IF | Dilution<br>WB | Reference | Provider |
| --- | --- | --- | --- | --- | --- |
| **Primary antibodies** |  |  |  |  |  |
| anti-ANXA1 | mouse monoclonal | 1: 200 | N/A | AMAb90558 | Atlas Antibodies,<br>Bromma, Sweden |
| anti-ANXA2 | rabbit polyclonal | 1: 25 | N/A | HPA046964 | Sigma-Aldrich, Saint-Quentin Fallavier, France |
| anti-ANXA8 | rabbit polyclonal | 1: 125 | N/A | SAB2700611 | Sigma-Aldrich, Saint-Quentin Fallavier, France |
| anti-CA4 | rabbit polyclonal | 1: 250 | N/A | HPA011089 | Sigma-Aldrich, Saint-Quentin Fallavier, France |
| anti-EDIL3 | rabbit polyclonal | 1: 500 | 1: 1, 000 | SAB2105802 | Sigma-Aldrich, Saint-Quentin Fallavier, France |
| anti-MFGE8 | rabbit polyclonal | N/A | 1: 1, 000 | non commercial | ProteoGenix,<br>Schiltigheim, France |
| anti-PDCD6IP | rabbit polyclonal | 1: 500 | N/A | HPA011905 | Sigma-Aldrich, Saint-Quentin Fallavier, France |
| **Secondary antibodies** |  |  |  |  |  |
| anti-rabbit<br>(IgG, H+L) | goat Alexa Fluor<br>488 | 1: 1, 000 | N/A | A11034 | ThermoFisher Scientific,<br>Waltham, Massachusetts |
| anti-mouse<br>IgG1 cross-adsorbed | goat Alexa Fluor<br>555 | 1: 1, 000 | N/A | A21127 | ThermoFisher Scientific,<br>Waltham, Massachusetts |
| anti-rabbit<br>(IgG, H+L) | goat Alexa Fluor<br>680 | N/A | 1: 20, 000 | A21065 | ThermoFisher Scientific,<br>Waltham, Massachusetts |

## FOOTNOTES

The abbreviations used are: ACC: amorphous calcium carbonate B: bone BSA: bovine serum albumin CaCO_3_: calcium carbonate Ca^2+^: calcium CO_2_: carbon dioxide D: duodenum EDS: energy-dispersive X-ray spectroscopy EELS: electron energy loss spectroscopy EV: extracellular vesicle HCO_3_^−^: bicarbonate K: kidney L: liver Ma: magnum Mya: million years ago O.C.T.: optimal cutting temperature compound PBS: phosphate buffered saline PCR: polymerase chain reaction p.o.: post-ovulation RI: red isthmus RT-PCR: reverse transcriptase polymerase chain reaction SAED: selected area electron diffraction SDS: sodium dodecyl sulfate SEM: scanning electron microscopy SEP: soluble eggshell proteins TBS: tris buffered saline TEM: transmission electron microscopy UF: uterine fluid Ut: uterus WI: white isthmus

